# Mechanism-based sirtuin enzyme activation

**DOI:** 10.1101/027243

**Authors:** Xiangying Guan, Alok Upadhyay, Sudipto Munshi, Raj Chakrabarti

**Author notes:** To whom correspondence should be addressed: Raj Chakrabarti, Ph.D. Division of Fundamental Research, ChakrabartiAdvanced Technology, 1288 Route 73 South, Mt. Laurel, NJ 08054, USA.

## Abstract

Sirtuin enzymes are NAD^+^-dependent protein deacylases that play a central role in the regulation of healthspan and lifespan in organisms ranging from yeast to mammals. There is intense interest in the activation of the seven mammalian sirtuins (SIRT1-7) in order to extend mammalian healthspan and lifespan. However, there is currently no understanding of how to design sirtuin-activating compounds beyond allosteric activators of SIRT1-catalyzed reactions that are limited to particular substrates. Moreover, across all families of enzymes, only a dozen or so distinct classes of non-natural small molecule activators have been characterized, with only four known modes of activation among them. None of these modes of activation are based on the unique catalytic reaction mechanisms of the target enzymes. Here, we report a general mode of sirtuin activation that is distinct from the known modes of enzyme activation. Based on the conserved mechanism of sirtuin-catalyzed deacylation reactions, we establish biophysical properties of small molecule modulators that can in principle result in enzyme activation for diverse sirtuins and substrates. Building upon this framework, we propose strategies for the identification, characterization and evolution of hits for mechanism-based enzyme activating compounds. We characterize several small molecules reported in the literature to activate sirtuins besides SIRT1, using a variety of biochemical and biophysical techniques including label-free and labeled kinetic and thermodynamic assays with multiple substrates and protocols for the identification of false positives. We provide evidence indicating that several of these small molecules reported in the published literature are false positives, and identify others as hit compounds for the design of compounds that can activate sirtuins through the proposed mechanism-based mode of action.

## Introduction

Sirtuin (silent information regulator) enzymes, which catalyze NAD^+^-dependent protein post-translational modifications, have emerged as critical regulators of many cellular pathways. In particular, these enzymes protect against age-related diseases and serve as key mediators of longevity in evolutionarily distant organismic models [1]. Sirtuins are NAD^+^-dependent lysine deacylases, requiring the cofactor NAD^+^ to cleave acyl groups from lysine side chains of their substrate proteins, and producing nicotinamide (NAM) as a by-product. A thorough understanding of sirtuin chemistry is not only of fundamental importance, but also of considerable medicinal importance, since there is enormous current interest in the development of new mechanism-based sirtuin modulators [2, 3]. The mechanism of sirtuin-catalyzed, NAD^+^-dependent protein deacylation is depicted in Fig. 1 [4-6].

**Figure 1.**
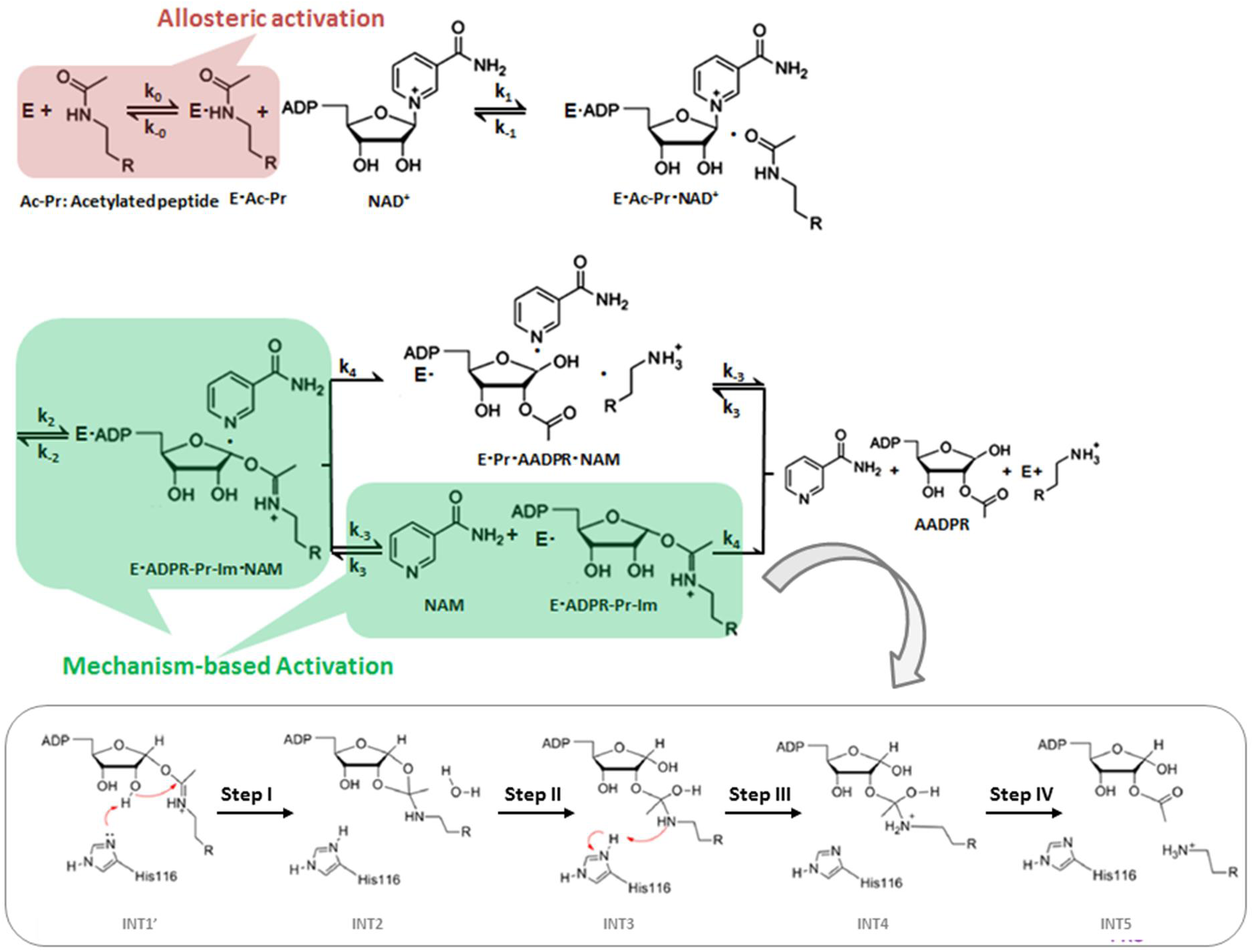
Chemical mechanism of sirtuin-catalyzed deacylation and modes of sirtuin activation. Following sequential binding of acylated peptide substrate and NAD^+^ cofactor, the reaction proceeds in two consecutive stages: i) cleavage of the nicotinamide moiety of NAD^+^ (ADP-ribosyl transfer) through the nucleophilic attack of the acetyl-Lys side chain of the protein substrate to form a positively charged O-alkylimidate intermediate, and ii) subsequent formation of deacylated peptide. For simplicity, all steps of stage ii as well as AADPR + Pr dissociation are depicted to occur together with rate limiting constant *k_4_*. **Red**: Allosteric activation increases the affinity of a limited set of peptide substrates for the SIRT1 enzyme only and requires an allosteric binding site. **Green**: Mechanism-based activation is a new mode of enzyme activation that relies on the conserved sirtuin reaction mechanism that are not limited to enhancement of affinity for selected peptide substrates. The schematic highlights mechanism-based activation through NAD^+^ K_m_ reduction rather than the K_d_ peptide reduction that known allosteric sirtuin activators elicit.

Recently, in order to extend mammalian healthspan and lifespan, intense interest has developed in the activation of the seven mammalian sirtuin enzymes (SIRT1-7). Prior work on sirtuin activation has relied exclusively on experimental screening, with an emphasis on allosteric activation of the SIRT1 enzyme. Indeed, small molecule allosteric activators of SIRT1 have been demonstrated to induce lifespan extension in model organisms such as mice [7, 8]. Allosteric activation is one of four known modes by which small molecules can activate enzymes [9]. Allosteric activators most commonly function by decreasing the dissociation constant for the substrate (the acylated protein dissociation constant *K_d,Ac_*_–Pr_ in the case of sirtuins).

Nearly all known sirtuin activators allosterically target SIRT1 and bind outside of the active site to an allosteric domain in SIRT1 that is not shared by SIRT2-7 [10]. Moreover, allosteric activators only work with a limited set of SIRT1 substrates [11, 12]. It is now known that other sirtuins -- including SIRT2, SIRT3 and SIRT6 -- and multiple protein substrates play significant roles in regulating mammalian longevity [13-15]. General strategies for the activation of any mammalian sirtuin (including activation of SIRT1 for other substrates) are hence of central importance, but not understood.

Aside from allosteric activation, enhancement of enzymatic activity via “derepression of inhibition” has been explored theoretically and experimentally. Theoretical models proposed have been limited to inhibitors that are exogenous to the reaction, and experimental studies have considered alleviation of product inhibition through competition with product binding [4]. These approaches can only enhance enzyme activity in the presence of inhibitor or product accumulation and hence are not included among the four known modes of enzyme activation.

Foundations for the rational design of mechanism-based sirtuin activators have been lacking, partly due to the absence of a clear understanding of the kinetics of sirtuin-catalyzed deacylation. Several types of mechanism-based sirtuin inhibitors have been reported recently in the literature, including Ex-527 and Sir-Real2 [16, 17]. However, mechanism-based activation has proven far more elusive, due to the difficulty in screening for the balance of properties needed for a modulator to have the net effect of accelerating catalytic turnover. While there are many ways to inhibit an enzyme’s mechanism, there are far fewer ways to activate it. These efforts have been hindered by the lack of a complete steady state kinetic model of sirtuin catalysis that accounts for the effects of both NAD^+^ and NAM on activity.

Small molecules that can activate sirtuins through modes of action other than the reduction of substrate K_d_ are of particular interest, because sirtuin activity often declines during aging for reasons other than substrate depletion. In the so-called “NAD^+^ world” picture of global metabolic regulation, the intracellular concentrations of the sirtuin cofactor NAD^+^, which can decrease with age, play a central role in regulating mammalian metabolism and health through sirtuin-dependent pathways [18]. Due to the comparatively high Michaelis constants for NAD^+^ (K_m,NAD+_’s) of mammalian sirtuins, their activities are sensitive to intracellular NAD^+^ levels [18]. The systemic decrease in NAD^+^ levels that accompanies organismic aging downregulates sirtuin activity and has been identified as a central factor leading to various types of age-related health decline [18], whereas increases in NAD^+^ levels can upregulate sirtuin activity and as a result mitigate or even reverse several aspects of this decline [18]. In addition to NAD+ cofactor concentration, the concentration of NAM is also believed to regulate sirtuin activity *in vivo*.

As such, NAD+ supplementation has emerged as a promising alternative to allosteric activation of sirtuins [18]. Unlike allosteric activators like resveratrol, which are SIRT1-specific and have not been successfully applied to other sirtuins [10], NAD^+^ supplementation can activate most mammalian sirtuins in a substrate-independent fashion. Moreover, allosteric activators cannot fully compensate for the reduction in sirtuin activity that occurs through NAD^+^ decline during aging. On the other hand, the effects of NAD^+^ supplementation are not specific to sirtuins and prohibitively high concentrations of NAD^+^, along with associated undesirable side effects, may be required to elicit the increases in sirtuin activity required to combat age-related diseases.

A preferred general strategy for activation of sirtuins (Fig. 1) would be to increase their sensitivity to NAD^+^ through a reduction of *K_m_*_,*NAD*+_. *K_m_*_,*NAD*+_ reduction would have a similar activating effect to NAD^+^ supplementation, but would be selective for sirtuins and could potentially even provide isoform specific sirtuin activation. Importantly, due to the sirtuin nicotinamide cleavage reaction that involves the NAD^+^ cofactor, modulation of *K_m_*_,*NAD*+_ may in principle be achievable by means other than altering the binding affinity of NAD^+^. Unlike allosteric activation that reduces *K_d,Ac_*_–Pr_, this approach could be applicable to multiplesirtuins and substrates.

Several compounds have been reported in the literature as being activators of sirtuins other than SIRT1 [19-22]. Concurrently, sirtuins whose preferred substrates are long chain acylated peptides were reported to have their activities on shorter chain acyl groups enhanced in the presence of certain fatty acids, further corroborating the feasibility of substrate-specific sirtuin activation [23]. However, some of the aforementioned compounds have not been characterized using initial rate assays [20,21] and others have been studied using only labeled activity assays that may be susceptible to false positives [21]. Moreover, studies that have employed label-free initial rate assays [22] have nonetheless not shed light on the underlying mechanism by which activation may occur. The theory presented in this paper, which is applied to one such compound as an example, may provide a unifying framework under which such compounds may be characterized and hits may be evolved into leads. Moreover, the experimental methods presented may help in uncovering false positive hits which may lead to inefficient usage of discovery research resources.

In this paper, we present a general framework for activation of sirtuin enzymes that is distinct from any of the known modes of enzyme activation, based on the fundamental mechanism of the sirtuin deacylation reaction. We first introduce a steady-state model of sirtuin-catalyzed deacylation reactions in the presence of NAD^+^ cofactor and endogenous inhibitor NAM, and then establish quantitatively how k_cat_/K_m,NAD+_ can be modified by small molecules, identifying the biophysical properties that small molecules must have to function as such mechanism-based activators. The principles introduced can also be generalized to the reduction of peptide substrate K_m_. through non-allosteric mechanisms. We propose strategies suitable for mechanism-based design of sirtuin activating compounds and characterize several small molecules that have been reported to be activators of sirtuins other than SIRT1 in order to determine whether these compounds possess the proposed characteristics of mechanism-based sirtuin activating compounds (MB-STACs) presented.

## Results

### Steady-state sirtuin kinetic modeling

To a greater extent than inhibitor design, rational activator design requires the use of a mechanistic model in the workflow. In this section we develop a steady state model for sirtuin-catalyzed deacylation that is suitable for a) investigation of the mode of action of mechanism-based sirtuin modulators, including activators; b) design of mechanism-based sirtuin activating compounds. We first summarize the state of knowledge regarding the sirtuin-catalyzed deacylation mechanism.

The sirtuin catalytic cycle (Fig. 1) is believed to proceed in two consecutive stages [4]. The initial stage (ADP-ribosylation) involves the cleavage of the nicotinamide moiety of NAD^+^ and the nucleophilic attack of the acyl-Lys side chain of the protein substrate to form a positively charged O-alkylimidate intermediate [4, 24]. Nicotinamide-induced reversal of the intermediate (the so-called base exchange reaction) causes reformation of NAD^+^ and acyl-Lys protein. The energetics of this reversible reaction affects both the potency of NAM inhibition of sirtuins and the Michaelis constant for NAD+ (K_m,NAD+_). The second stage of sirtuin catalysis, which includes the rate-determining step, involves four successive steps that culminate in deacylation of the Lys side chain of the protein substrate and the formation of O-acetyl ADP ribose coproduct [4, 6, 25].

A tractable steady state model suitable for the purpose of mechanism-based sirtuin activator design must account for the following important features:

- The calculated free energy of activation for nicotinamide cleavage (ADP -ribosylation of the acyl-Lys substrate) in the bacterial sirtuin enzyme Sir2Tm as computed through mixed quantum/molecular mechanics (QM/MM) methods is 15.7 kcal mol^-1^ [5, 26]. An experimental value of 16.4 kcal mol^-1^ for the activation barrier in the yeast sirtuin homolog Hst2 was estimated from the reaction rate 6.7 s^-1^ of nicotinamide formation. The nicotinamide cleavage reaction is endothermic, with a computed Δ*G* of 4.98 kcal mol^-1^ in Sir2Tm [26].
- The calculated free energy of activation for the rate limiting chemistry step (collapse of the bicyclic intermediate) from QM/MM simulations is 19.2 kcal mol^−1^ for Sir2Tm [27], in good agreement with the experimental value of 18.6 kcal/mol^-1^ estimated from the *k*_cat_ value of 0.170 ± 0.006 s^-1^ [28] (0.2 +/− 0.03 s^-1^ for Hst2 [24]). We note that the relative magnitudes of the rate constants for the two slowest chemistry steps may vary for other sirtuins, like mammalian sirtuins. For some sirtuins, product release may be rate limitng.
- The remaining steps in the catalytic cycle are significantly faster than the above steps. The other chemistry steps in stage 2 of the reaction are effectively irreversible [27], as is product release in the presence of saturating peptide concentrations.

We hence include in our kinetic model representations of all steps in stage 1 of the reaction, including the nicotinamide cleavage/base exchange and nicotinamide binding steps. However, for simplicity, we do not include in the present model a representation of each of the individual chemistry steps in stage 2 of the reaction or final product release, instead subsuming these steps under the smallest rate constant, which we call k_cat_. Since all these steps are effectively irreversible, the full steady state model including these steps can be immediately derived from the basic model through simple modifications, to be described in a subsequent revision, that are not essential to the analysis of mechanism-based activation. The above observations motivate the kinetic model represented in Fig. 2 [29]. This Figure shows a general reaction scheme for sirtuin deacylation including base exchange inhibition.

**Figure 2.**
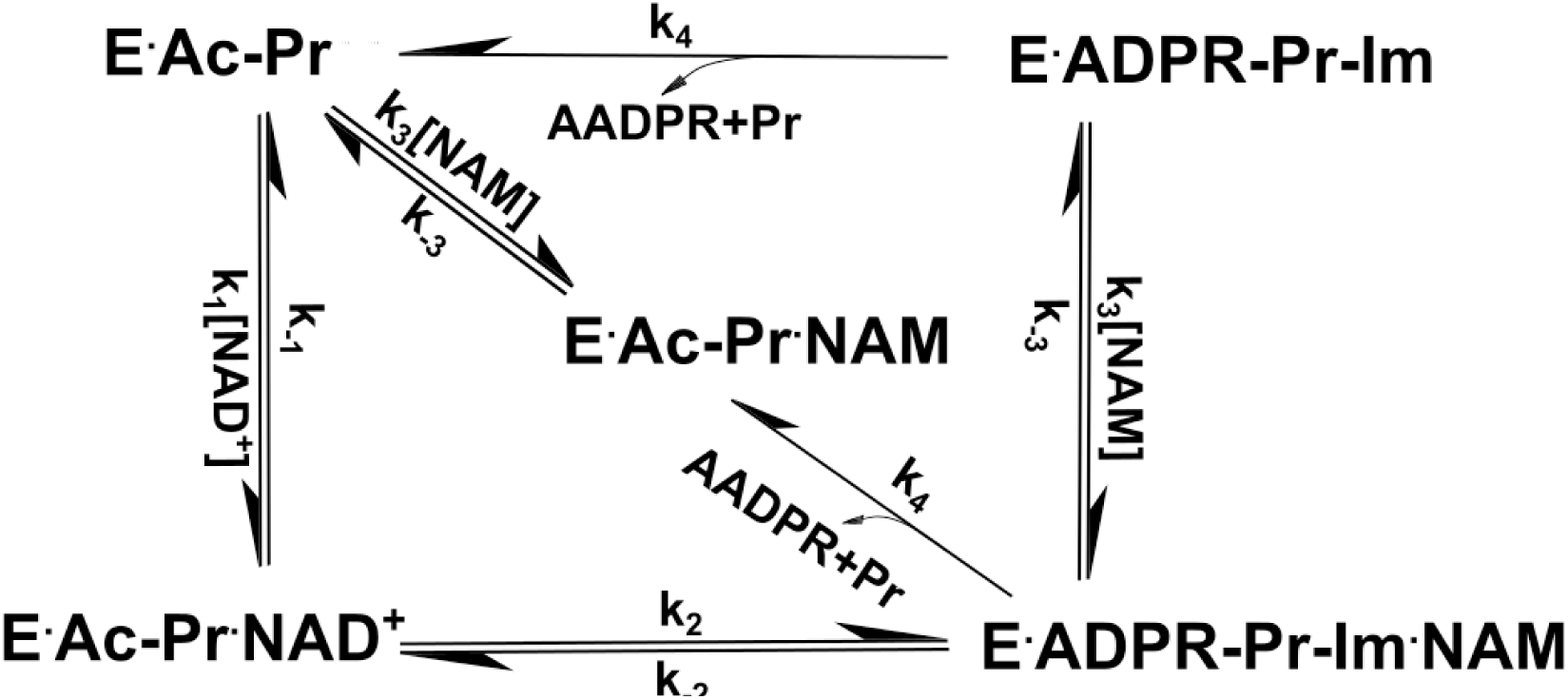
General model for sirtuin-catalyzed deacylation in the presence of NAD^+^ and NAM. This model, based on the reaction mechanism depicted in Fig. 1, provides a minimal kinetic model that captures the essential features of sirtuin deacylation kinetics suitable for predicting the effects of mechanism-based modulators on sirtuin activity. In the presence of saturating Ac-Pr, E is rapidly converted into E. Ac-Pr and NAM binding to E can be neglected, resulting in a simplified reaction network with 5 species. Ac-Pr, acetylated peptide; ADPR, adenosine diphosphate ribose; AADPR, O-acetyl adenosine diphosphate ribose.

Because of the physiological benefits of improving catalytic efficiency under NAD depletion rather than peptide depletion conditions, we assume saturating peptide conditions in our kinetic modeling in this paper. However, precisely analogous equations could be derived for saturating NAD, in the event that activation under peptide depletion conditions is desired.

The reaction mechanism of sirtuins precludes the use of rapid equilibrium methods for the derivation of even an approximate initial rate model; steady-state modeling is essential. The rate equations for the reaction network in Fig. 2 enable the derivation of steady-state conditions for the reaction. Solving the linear system of algebraic steady-state equations and mass balance constraints for the concentrations

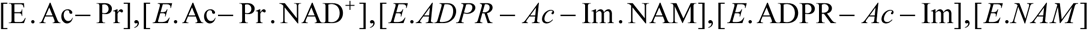
in terms of the rate constants and [NAD+],[NAM], which are assumed to be in significant excess and hence approximately equal to their initial concentrations [NAD+]_0_,[NAM]_0_ respectively, we obtain expressions of the form (1):

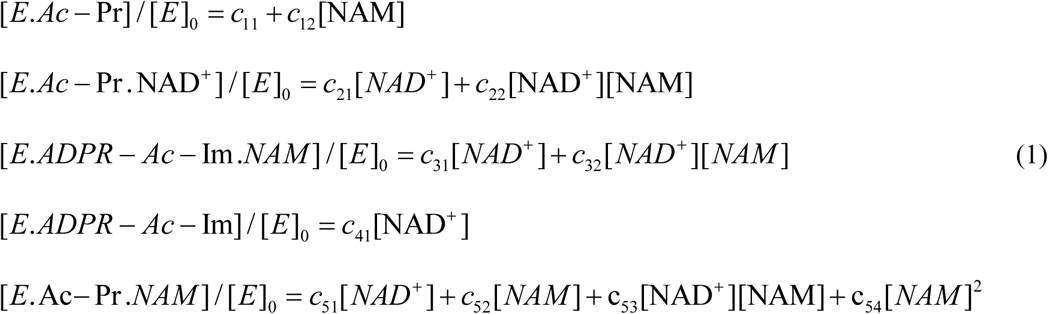
where the term c_54_ that is second order in [NAM] will be omitted from the analysis below. Expressions for the c_ij_’sare provided in the Appendix. The initial rate of deacylation can be then expressed

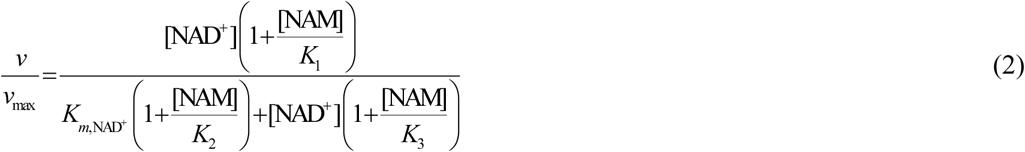
with

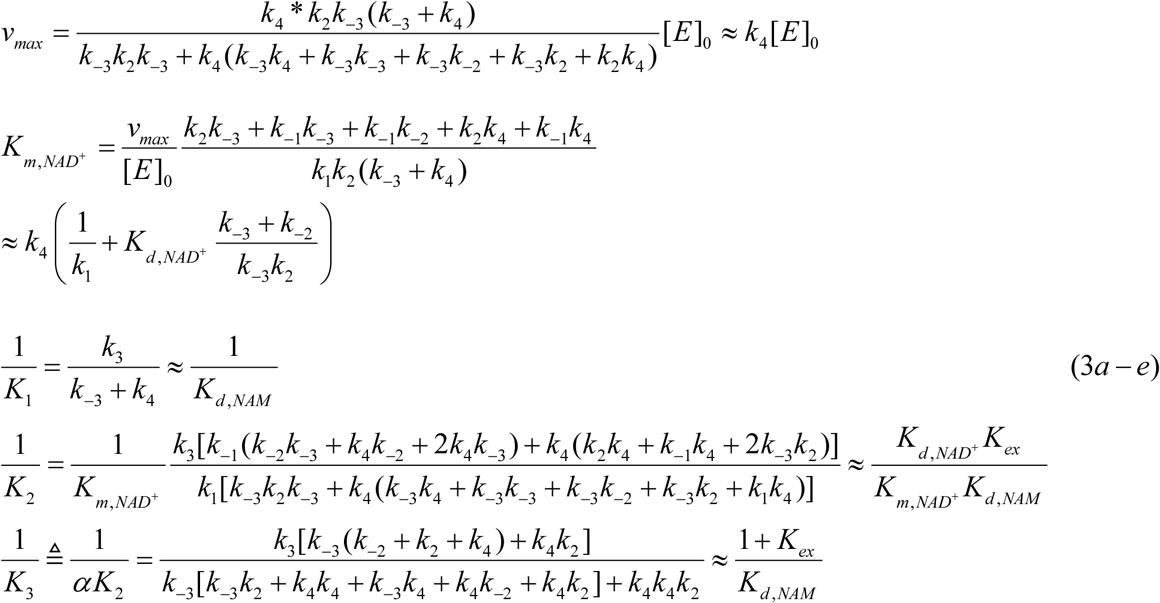
where K_ex_ ≡ k_-2_/k_2_ and the approximations refer to the case where *k*_4_ ≪ *k_j_*, *j* ≠ 4. The quality of this approximation can be assessed for the chemistry steps based on QM/MM simulation data, which was cited above for yeast and bacterial sirtuins, or from experimental methods for the estimation of all rate constants in the model (see below). Expression (2) can be used to calculate the initial rate of sirtuin-catalyzed deacylation for specified intracellular concentrations of NAD+ and NAM, assuming the rate constants are known.

Equation (2) is typically represented graphically in terms of either double reciprocal plots at constant [NAM] or Dixon plots at constant [NAD+]. In the former case, the slope of the plot (1/v vs 1/[NAD+]) at [NAM]=0 is 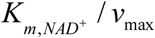, for which the expression is:

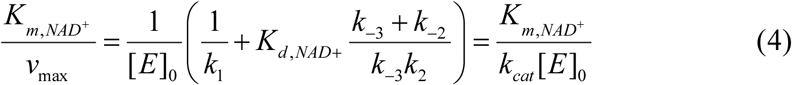
whereas for the Dixon plot, the expression for the slope at 1/[NAD+]=0 is approximately [29]:

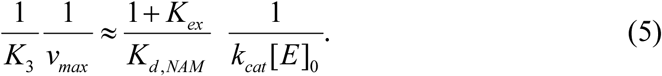

We note that equation (4) for catalytic efficiency applies irrespective of the small k_4_ approximation.

The steady state parameter *α* in equation (3e), which is a measure of the extent of competitive inhibition by the endogeneous inhibitor NAM against the cofactor NAD+, can be expressed in terms of the ratio of K_d,NAD+_ and K_m,NAD+_ [29]:

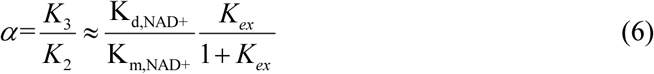
which, together with expression (3b) for K_m,NAD+_, demonstrates how the kinetics of inhibition of deacylation by NAM can reveal differences in NAD^+^ binding affinity and nicotinamide cleavage rates among sirtuins. The origins of different NAD+ binding affinities among sirtuins were studied structurally and computationally in [29]. Given that K_ex_ is generally >> 1 for sirtuins, it is apparent from eqn (6) that the difference in magnitudes of K_d,NAD+_ and K_m,NAD+_ for sirtuins is captured by *α* for sirtuins satisfying the above approximations. K_m,NAD+_, not K_d,NAD+_ alone, determines the sensitivity of sirtuin activity to NAD+, and can vary across this family of enzymes.

In addition to the approximation *k*_4_ ≪ *k_j_*, *j* ≠ 4, several experimental observations can further simplify the form of the expressions (4) for the sirtuin steady state constants. First, we assume *k*_−3_ ≫ *k_j_*, *j* ≠ −2 based on viscosity measurements that suggest NAM dissociates rapidly following cleavage [30]. Under this approximation, the expression for K_m,NAD+_ becomes:

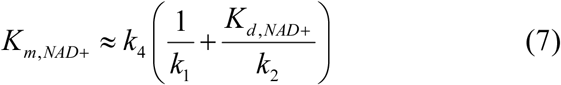

Such approximations will be studied in greater detail in a subsequent work.

As can be seen from eqn (3b), the kinetics of the nicotinamide cleavage reaction and the rate limiting step of deacylation both play essential roles in determining the value of K_m,NAD+_. Note that in rapid equilibrium models of enzyme kinetics, which are not applicable to sirtuins, K_m_ ≈ K*_d_*. The difference between K_d,NAD+_ and K_m,NAD+_ has important implications for mechanism-based activation of sirtuins by small molecules [29]. In particular, as we will show in this work, decrease of K_m,NAD+_ independently of K_d,NAD+_ can increase the activity of sirtuins at [NAM]=0. The kinetic model above establishes foundations for how this can be done.

### Mechanism-based sirtuin activation

A prerequisite for enzyme activation is that the modulator must co-bind with substrates – NAD^+^ and acylated peptide in the case of sirtuins. Within the context of enzyme inhibition, two modes of action display this property: noncompetitive and uncompetitive inhibition. Noncompetitive inhibitors bind with similar affinities to the apoenzyme and enzyme-substrate, enzyme-intermediate or enzyme-product complexes whereas uncompetitive inhibitors bind with significantly lower affinity to the apoenzyme. Traditional noncompetitive inhibitors reduce v_max_ and increase K_m_ to a much smaller extent. On the other hand, uncompetitive inhibitors decrease substrate K_m_ as a result of their preferential binding, but decrease v_max_ by at most the same amount, thus leading to no net increase (generally, a decrease) in catalytic efficiency. Both are specific examples of the more general notion of a mixed noncompetitive modulator that co-binds with substrates. In the case of sirtuins, examples of noncompetitive inhibitors include SirReal2 [18], whereas examples of uncompetitive inhibitors include Ex-527 [17]. Though some known sirtuin inhibitors may satisfy the requirement of cobinding with substrates, they do not possess other critical attributes necessary for mechanism-based enzyme activation. While such compounds may have promising properties as potential hits for the development of mechanism-based activators, prior studies have only characterized their kinetic effects in terms of traditional rapid equilibrium formulations of enzyme inhibition, rather than a steady-state formulation for mechanism-based enzyme modulation. Moreover, as noted above, several compounds have recently been reported to activate sirtuins other than SIRT1, but none of these have their mechanisms characterized within an appropriate theoretical framework.

Previous attempts to develop a general approach to sirtuin activation [30, 31] only considered competitive inhibitors of base exchange, which cannot activate in the absence of NAM. This is not actually a form of enzyme activation, but rather derepression of inhibition. Such derepression modalities based on competitive inhibition of product binding cannot be hits for activator design, since these compounds or their relatives cannot cobind with substrates. By contrast, here we present paradigms and design criteria for activation of sirtuins in either the absence or presence of NAM. Based on expression (3b) for K_m,NAD+_, it is in principle possible to activate sirtuins (not just SIRT1) for any substrate by alteration of rate constants in the reaction mechanism other than k_1_,k_-1_ and k_cat_, so as to reduce K_m,NAD+,_ not only K_d,Ac-Pr_ as with allosteric activators, which increase the peptide binding affinity of SIRT1 in a substrate-dependent fashion. We now explore how this may be achieved by augmenting the kinetic model to include putative mechanism-based activators (A) that can bind simultaneously with NAD+ and NAM. Fig. 3 depicts the reaction diagram for mechanism-based activation of sirtuins. Note that only the top and front faces of this cube are relevant to the mechanism of action of the previously proposed competitive inhibitors of base exchange [30, 31].

**Figure 3.**
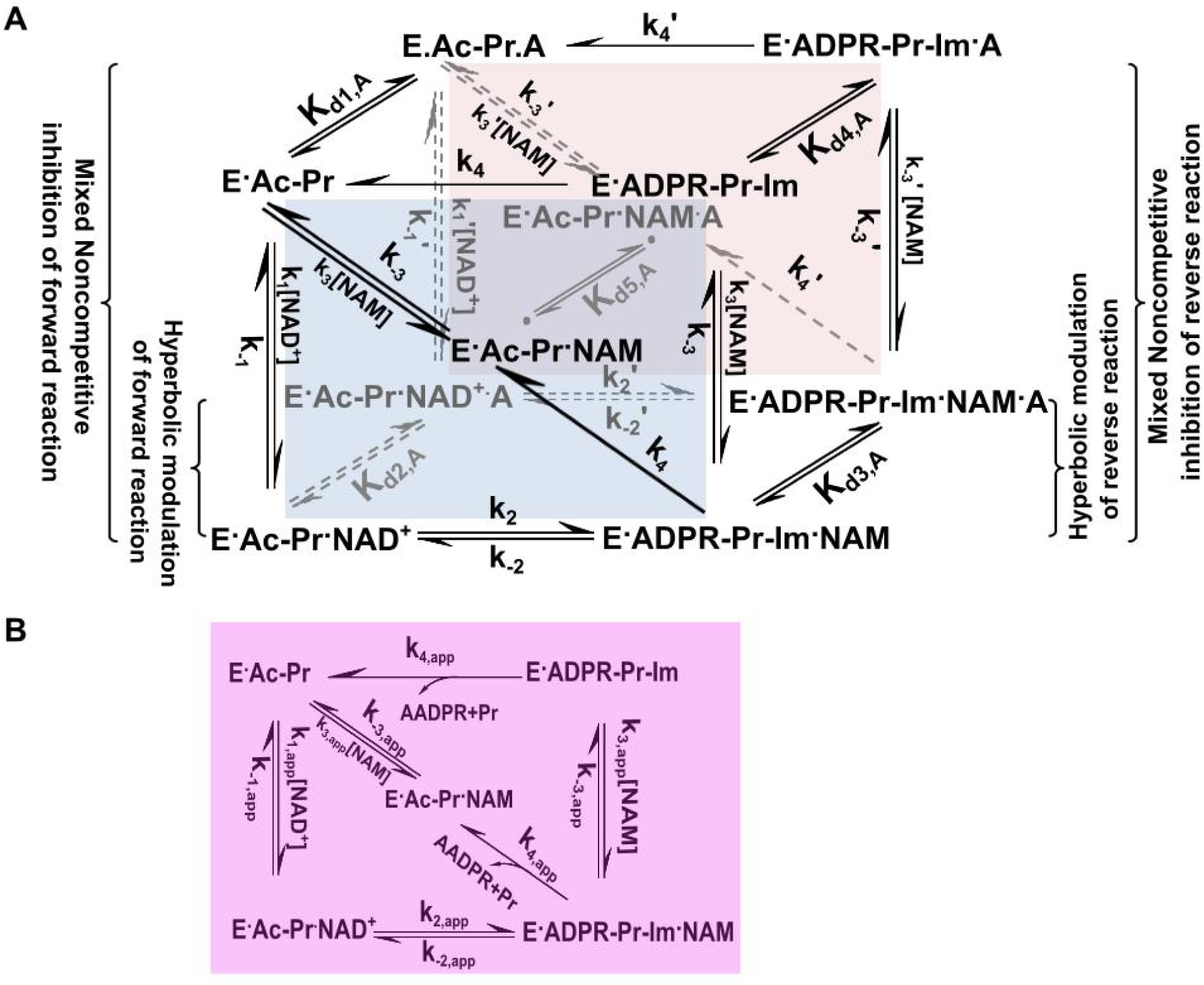
General model for mechanism-based sirtuin enzyme activation. **(A)** The front face of the cube (blue) depicts the salient steps of the sirtuin reaction network in the absence of bound modulator. The back face of the cube (red) depicts the reaction network in the presence of bound modulator (denoted by “A”). Each rate constant depicted on the front face has an associated modulated value on the back face, designated with a prime, that is a consequence of modulator binding. **(B)** The purple face is the apparent reaction network in the presence of a un-saturating concentration of modulator.

At any [A] there exist apparent values of each of the rate constants in the sirtuin reaction mechanism. These are denoted by “app” in the Figure. There are also corresponding “app” values of the steady state, Michaelis, and dissociation constants in equation (3). Moreover, at saturating [A] of a known activator, the modulated equilibrium and dissociation constants (which do not depend on determination of steady state species concentrations) can be estimated with only deacylation experiments according to the theory presented above. The exchange equilibrium constant *K_ex_* ′ and NAD^+^,NAM dissociation constants *K_d,NAD+_* ′ *K_d_*_,*NAM*_ ′ in the presence of A are related to their original values as follows:

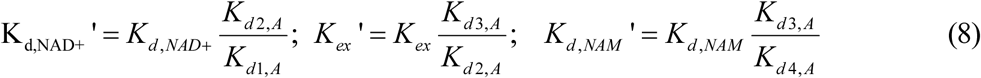
where the *K_d_*_,*A*_’s are the dissociation constants for A depicted in Fig. 3.

In order to predict the effect on 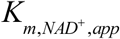 of a modulator with specified relative binding affinities for the complexes in the sirtuin reaction mechanism, it is important to develop a model that is capable of quantifying, under suitable approximations, the effect of such a modulator on the apparent steady state parameters of the enzyme. Since the full steady state expression relating the original to the apparent rate constants has many terms containing products of additional side and back face rate constants, we use a rapid equilibrium segments approach to arrive at simple definitions of the apparent Michaelis constant and other steady state constants for the reaction in terms of the original expressions for these constants and the dissociation constants for binding of A to the various complexes in the sirtuin reaction mechanism. This provides a minimal model with the least number of additional parameters required to model sirtuin activation mechanisms. In our treatment, we will assume that rapid equilibrium applies on both the side faces and the back face. Under this approximation, at low [A] the expressions for the induced changes in each of the rate constant products appearing in the coefficients c_ij_ and c_i’j’_, i’=i of equation (1) (see Appendix for expressions for these products) are the same and linear in [A]. For example, in the case of *E*.*Ac* − Pr, the steady state species concentrations become:

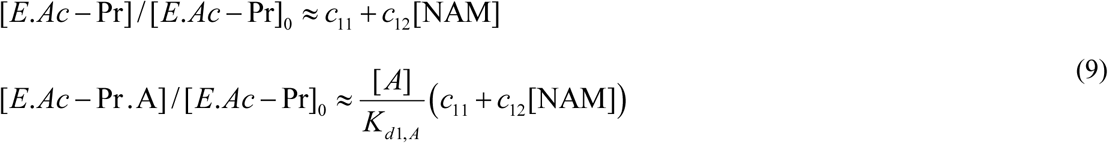

The rapid equilibrium segments expressions for all species concentrations in equation (1) in the presence of A are provided in the Appendix.

Expressions for apparent values of all steady state parameters introduced in equation (3) (i.e., modulated versions of constants *v*_max_, *K_m,NAD_*_+,_*K*_1_, *K*_2,_ *K*_3_) in the presence of a given [A] can now be derived. In the following, several types of approximations will be invoked:

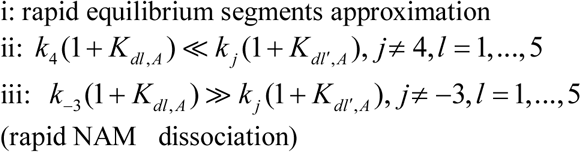

In particular, we emphasize that assumption (ii) may not hold for several mammalian sirtuins, but we apply this approximation to simplify the equations and provide physical insight. The general equations without this approximation can readily be derived using the principles introduced.

- 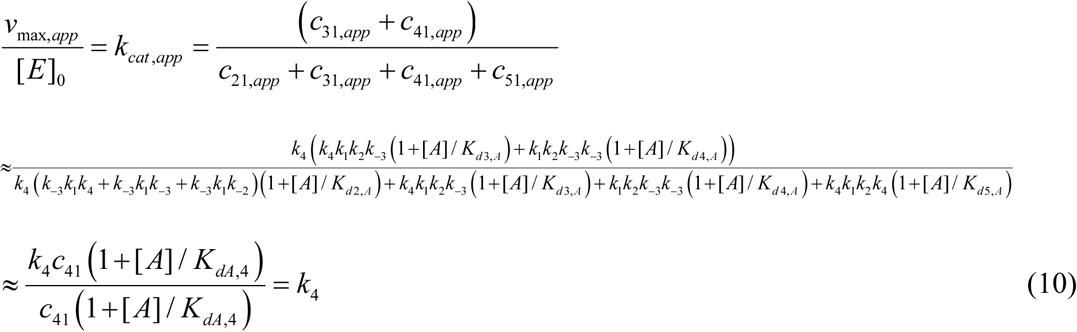
- 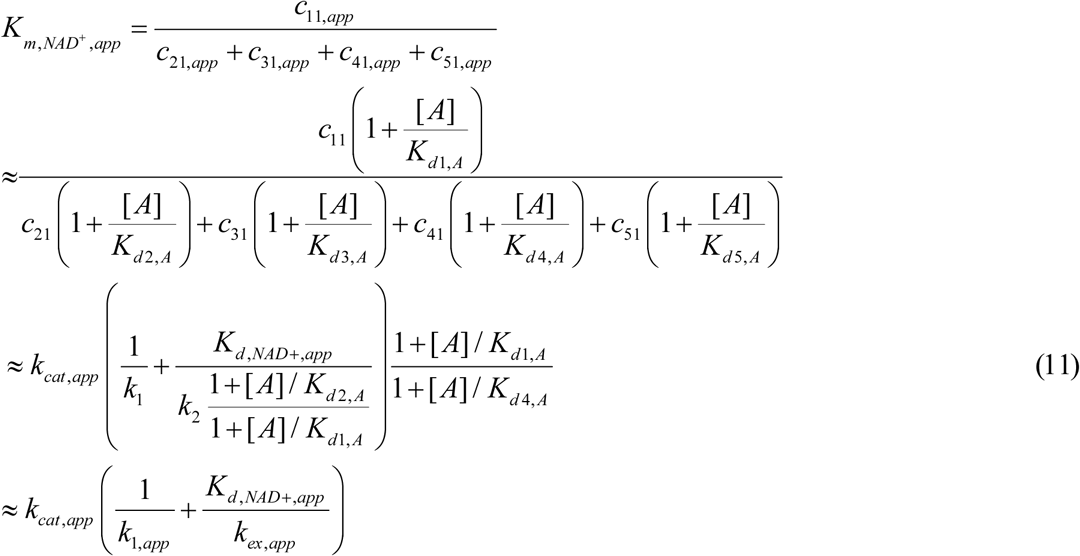
- 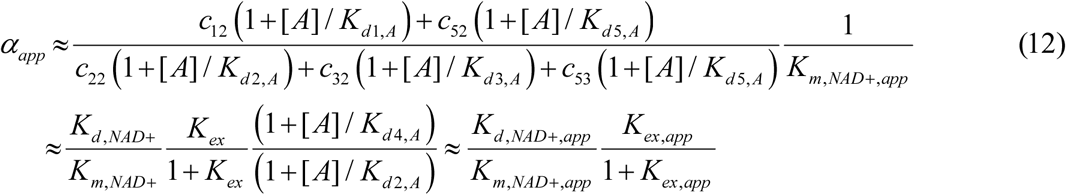
- 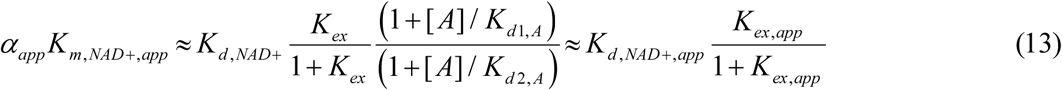
- Under approximation (ii), *K*_3_ isolates nicotinamide cleavage / base exchange-specific effects.

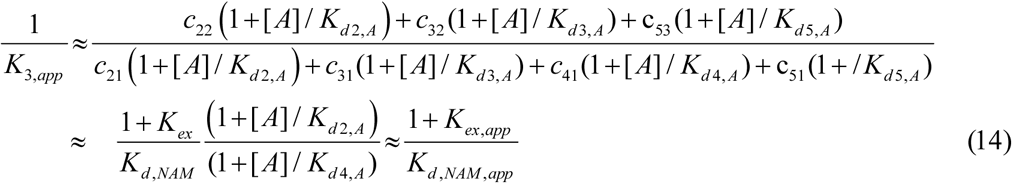
- 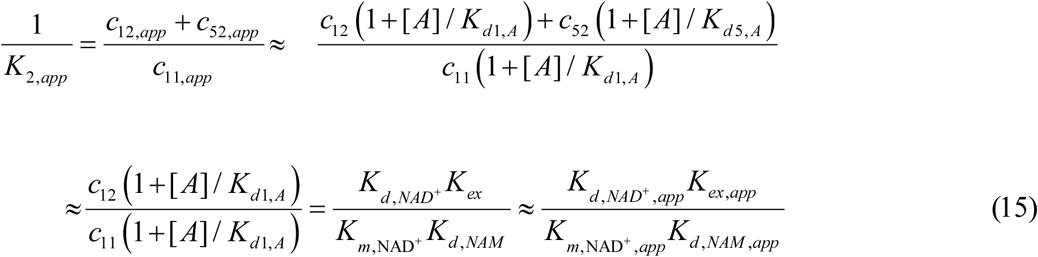

Regarding the quality of the approximations in this case, note from (15) and (A1) that unlike any of the other steady-state parameters, the modulation 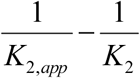 induced by [A] is proportional to *k_cat_* under the rapid equilibrium segments approximation (first approximation above). Hence, if one is interested in estimating the sign of this modulation, the small *k_cat_* approximation (second approximation above) should not be applied. Also, under the rapid equilibrium segments approximation, *K*_2,*app*_ is the only constant that relies on a ratio of two *c*_ij_’*s* with *i* ′*= i*, *j* ′ ≠ *j*, and hence the ratio of the same factor in [A].

- 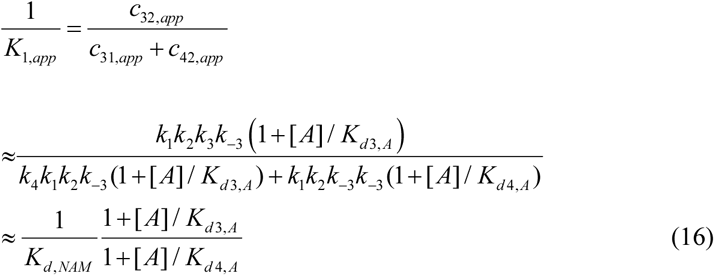

### Conditions for mechanism-based activation

We now consider thermodynamic conditions on the binding of a modulator A for mechanism-based sirtuin activation under the rapid equilibrium segments approximation, along with the expected changes in the steady state, equilibrium and dissociation constants in the sirtuin reaction mechanism.

First, according to equation (10), *v*_max_/[*E*]_0_ is roughly unchanged within this family of mechanisms as long as the *K_d,A_*’s for [A] binding to the various represented complexes in the reaction mechanism satisfy condition (iii). This is reasonable as long as the modulator does not lead to a significant increase in coproduct binding affinity and reduction in coproduct dissociation rate (for example, the stabilization of a closed loop conformation). As in the case of Ex-527 [17], the latter can render product dissociation rate limiting and reduce k_cat_,. If k_cat_ is reduced by the modulator, the net extent of activation will be reduced. However, even if this is the case, reduction in k_cat,app_ will not affect catalytic efficiency k_cat,app_/K_m,app_. Hence it is justified to omit binding of A to the coproduct complex from the mechanism-based activation model. Moreover, binding of A will not substantially reduce k_cat,app_ if the rate limiting chemistry step is much slower than product release.

The goal of mechanism-based activation is to increase k_cat,app_/K_m,app._. According to equation (11),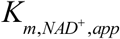 will be smaller than 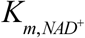 if *K_d_*_1,*A*_/*K_d_*_4,*A*_ ≥ (*K_d_*_1,*A*_/*K_d_*_2,*A*_)(*K_d_*_2,*A*_/*K_d_*_3,*A*_)(*K_d_*_3,*A*_/*K_d_*_4,*A*_) >1. To identify mechanisms by which this can occur in terms of the steps in the sirtuin-catalyzed reaction, we consider in turn each of these three respective ratios of *K_d_*_,*A*_’s (or equivalently, the ΔΔ*G* ‘s of the NAD^+^ binding, exchange, and NAM binding reactions as implied by equation (8)) induced by A binding.

According to equation (13), *K*_d1,A_/*K*_d2,A_ < 1 would imply that A binding increases the binding affinity of NAD^+^ to the E.Ac-Pr complex. This is the primary means by which allosteric activators enhance activity, but not the only possibility for mechanism-based activation. In principle, it is possible for a mechanism-based activator to reduce K_m,NAD+_even if K_d1,A_ ≥ K_d2,A_. In this case, in order to have 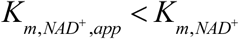, we require (*K_d_*_2,*A*_/*K_d_*_3,*A*_)(*K_d_*_3,*A*_/*K_d_*_4,*A*_) > *K_d_*_1,*A*_/*K_d_*_2,*A*_ or equivalently, according to (8), (*K_d_*_,*NAM*_′/*K_ex_*′)(*K_ex_*/*K_d_*_,*NAM*_) > *K_d_*_,*NAD*+_′/*K_d_*_,*NAD*+_. The decrease in 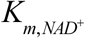 can then be due to modulation of the exchange rate constants that induces a decrease in *K_ex_*, an increase in *K_d,NAM_*, or both. If *K_d,NAM_* changes in the presence of modulator, this corresponds to mixed noncompetitive inhibition [29] of base exchange (Fig. 3).

As we have previously shown [29], the nicotinamide moiety of NAD^+^ engages in nearly identical interactions with the enzyme before and after bond cleavage. The salient difference is a conformational change in a conserved phenylalanine side chain (e.g., Phe33 in Sir2Tm, Phe157 in SIRT3) that destabilizes NAM binding after bond cleavage [32, 33]. Since NAM binding is already destabilized by the native protein conformation in this way, and since under rapid NAM dissociation (approximation iii above, which is believed to hold for sirtuins [23]) k_2_ and k_-2_ do not appear in the expression for K_m_, 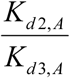 is likely to make the dominant contribution to 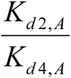 for activators Note that there is ample scope for modulation of Δ*G_ex_* by the modulator due to the coupling of the endothermic nicotinamide cleavage/ADP ribosylation reaction (exothermic base exchange reaction) to a conformational change in the sirtuin cofactor binding loop. For example, Δ*G_ex_* for Sir2Tm has been calculated to be -4.98 kcal/mol [26]. For comparison, Δ*G_bind,NAM_* for Sir2Af2 was estimated to be -4.1 kcal/mol [30] and Δ*G_bind,NAM_* for SIRT3 was estimated to be <= -3.2 kcal/mol [29]. Although the value of *K_d_*_2_/*K_d_*_4_ required for activation is likely to be achieved primarily by altering the free energy change of the nicotinamide cleavage reaction, our model accommodates the possibility of arbitrary combinations of ΔΔ*G_ex_* and ΔΔ*G_bind_*_,*NAM*_.

In our original model for sirtuin kinetics in Fig. 3, we assumed that both *K_d_*_,*NAM*_’s–namely, those for dissociation of NAM from E.Ac-Pr.NAM and E.ADPR-Pr-Im.NAM – are roughly equal. We maintain this condition in the presence of A binding, which is reasonable given that A is assumed to not interact directly with the peptide or ADPR moiety. Hence, we have:

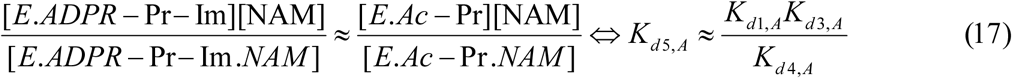

Returning to equation (11) for 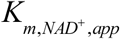 and substituting (1+[*A*]/*K_d_*_2,*A*,_)/(1+[*A*]/*K_d_*_1,*A*_) ≥ 1, the rapid equilibrium assumptions applied to the present system imply that in order to activate the enzyme at [NAM]=0, if A does not improve cofactor binding affinity it must increase k_1_ (k_1,app_>k_1_), k_2_ (k_2,app_>k_2_) or both. The rapid equilibrium segments model is not able to distinguish between these scenarios. If A increases 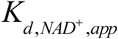, it is unlikely that an increase k_1_ will achieve activation. An increase in k_2_ implies acceleration of the rate of nicotinamide cleavage. In the rapid equilibrium segments framework, this occurs through preferential stabilization of the E.ADPR-Pr-Im complex. Note that across all sirtuins studied, nicotinamide cleavage induces structural changes (for example, unwinding of a helical segment in the flexible cofactor binding loop [34, 35]) and such changes might enable preferential stabilization of the E.ADPR-Pr-Im complex, in a manner similar to the stabilization of specific loop conformations by mechanism-based inhibitors [17]. Indeed, stabilization of alternative, non-native conformations of this loop have been observed crystallographically by reported activators of sirtuins other than SIRT3, including both long-chain fatty acids [23] and recently reported activators of SIRT5 and SIRT6 [22]. We discuss below the biophysical underpinnings whereby an increase in a forward rate constant could be achieved through preferential stabilization of the intermediate complex.

Given that an open loop conformation favors NAD+ binding, if a closed loop conformation is stabilized by the modulator, we expect the following thermodynamic conditions on the binding of A to the various complexes in the sirtuin reaction mechanism:

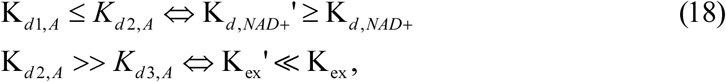
where the >> sign signifies that *K_d_*_2,*A*_/*K_d_*_3,*A*_ > *K_d_*_1,*A*_/*K_d_*_2,*A*_. We also require the necessary but not sufficient condition that *K_d_*_2,*A*_/*K_d_*_4,*A*_ > *K_d_*_1,*A*_/*K_d_*_2,*A*_. As noted above, increasing *K_d_*_3,*A*_/*K_d_*_4,*A*_ (which would destabilize NAM binding) is not considered as a mode of activation since NAM is believed to already dissociate quickly from the native active sites of sirtuins. Within conventional nomenclature, decrease in K_ex_ corresponds to hyperbolic (or partial) noncompetitive inhibition [29] of base exchange/activation of nicotinamide cleavage (as opposed to complete quenching of the base exchange reaction; see Fig. 3).

We now consider the effects of binding of such a modulator A that favors a closed cofactor loop conformation on the remaining steady state constants.

- *α_app_*: According to equation (12), the aforementioned condition that *K_d_*_2,*A*_/*K_d_*_4,*A*_ ≥ *K_d_*_1,*A*_/*K_d_*_2,*A*_ implies an increase in *α*.
- *K*_2,*app*_: With conditions (18), equation (14) predicts a limited change in *K*_2_.
- *K*_3,*app*_: According to equation (14), in the presence of such a mechanism-based activator, *K*_3_ is expected to increase by a factor more than that for *K*_2_ under the rapid equilibrium segments, approximation. This can occur due to an increase in *K_d_*_,*NAM*_ a decrease in K_ex_, or both.
- *K*_1,*app*_: Reduction in *K*_1_ through a reduction in *K_d_*_3,*A*_/*K_d_*_4,*A*_ (eqn 16) can increase K_m,NAD+_. A modulator has more favorable properties if *K*_1_ does not decrease significantly, but as discussed above, an increase in *K*_1_ will generally not be sufficiently for activation. Moreover, if *K*_3_ increases in the presence of modulator, a decrease in *K*_1_ may not be consequential.

Fig. 4 depicts the model-predicted changes to the various steady state, Michaelis and dissociation constants in the sirtuin reaction mechanism in the presence of such a modulator.

**Figure 4.**
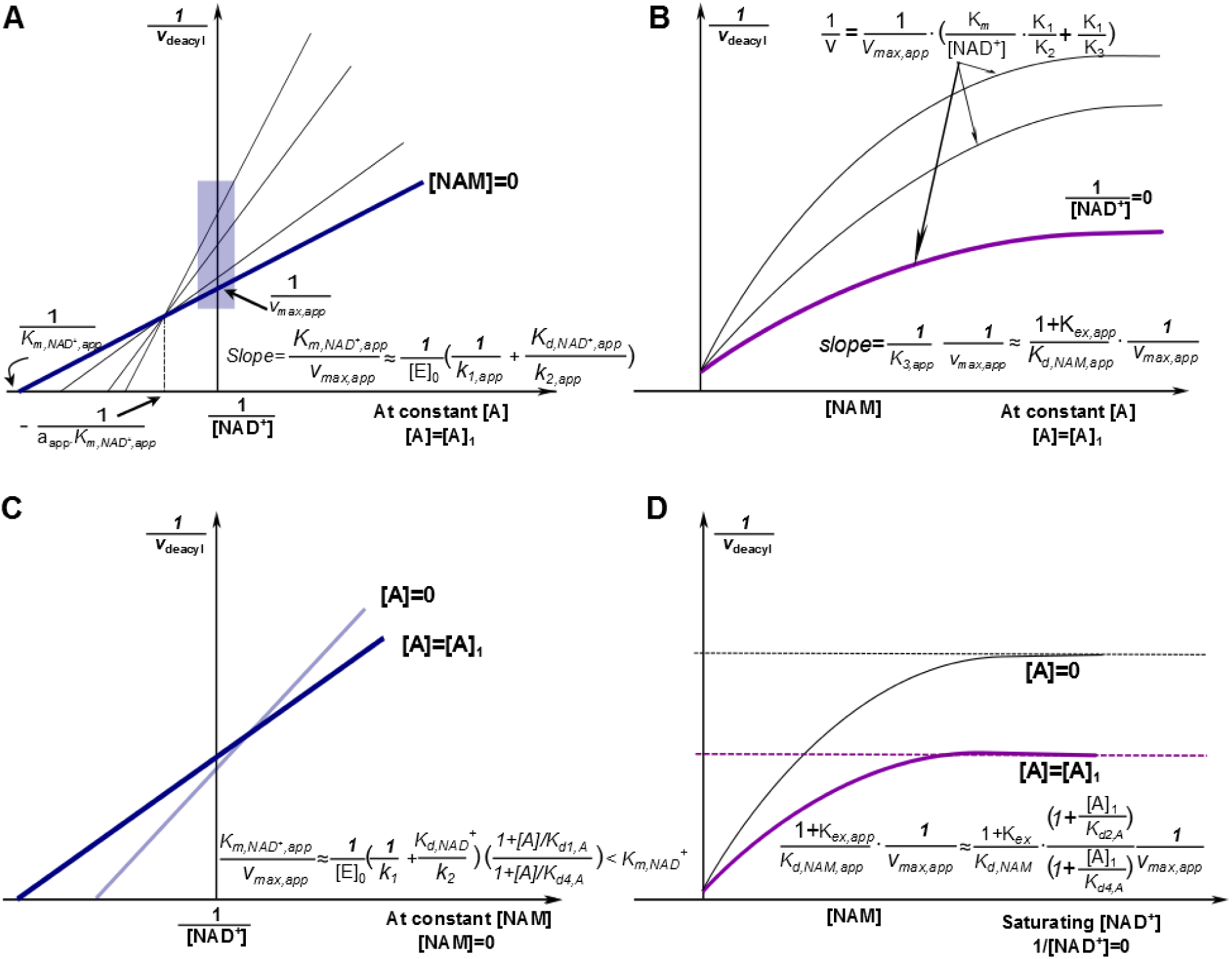
Mechanism-based activation of sirtuin enzymes: predicted steady-state properties and dose-response behavior. (a) Double reciprocal plots for deacylation initial rate measurements in the presence of activator. The blue box on the y-axis highlights the data that is used to construct the Dixon plot at saturating [*NAD^+^*] depicted in (b). (b) Dixon plots for deacylation initial rate measurements in the presence of activator. The arrows point to predicted plateaus in these curves. (c) Comparison of double reciprocal plots at [NAM] = 0 uM in the presence and absence of activator. (d) Comparison of Dixon plots at 1/[NAD^+^] = 0 in the presence and absence of activator. “A” denotes a mechanism-based sirtuin activating compound. Note that the model depicted omits the term quadratic in [NAM] in eqn (1) and the plateaus/dotted lines shown in the Dixon plots are the asymptotic values to which the model-predicted rates converge in the absence of this term.

**Figure 5.**
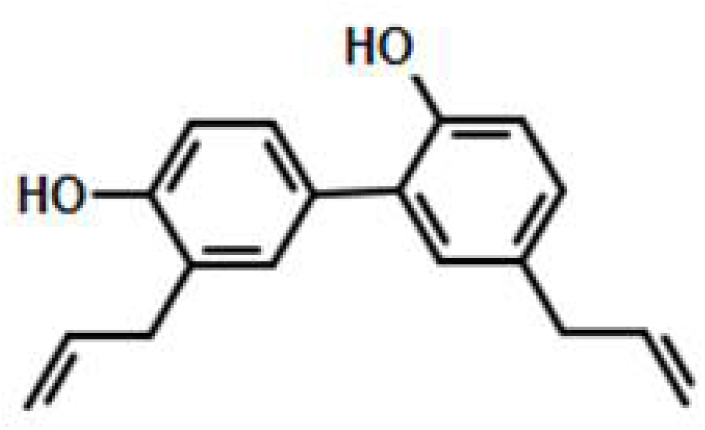
Structure of honokiol (HKL), 5, 3′-Diallyl-2, 4′-dihydroxybiphenyl (Honokiol).

The conditions for activation described above do not need to hold for a hit compound for mechanism-based activation. A hit compound may be defined as one that satisfies a subset of the conditions enumerated above, and may also display comparatively little inhibition in the pre-steady state burst phase. Such compounds may be capable of undergoing further improvement for substrate-specific activation of sirtuins like SIRT3, under physiologically relevant NAD^+^ depletion conditions. For example, it is possible that catalytic efficiency does not increase in the absence of NAM, but does so in its presence (for example, due to increase in the K_3_ parameter above). Note that due to nonzero physiological concentration of NAM in the cell, reduction of NAM inhibition can also contribute to activation under physiologically relevant conditions. Alternatively, the relative rates of deacylation in the presence and absence of modulator could converge under certain combinations of [NAD^+^] and [NAM].

### Potential means of increasing k_2_

From the standpoint of chemical mechanisms of activation, the theory presented raises the important question of how the nicotinamide cleavage rate k_2_ of sirtuins can be accelerated by a ligand that binds to the various complexes in the deacylation reaction with the specified relative affinities, as predicted by equation (11). It is important to note in this regard that the nicotinamide cleavage reaction in sirtuins is generally believed to be endothermic, which enables effective NAM inhibition of the reaction [26, 36]. Unlike exothermic reactions, stabilization of products in endothermic reactions can decrease the activation barrier for the forward reaction, due to the fact that the transition state resembles the products more than the reactants. The energetics of this reaction, including the role of protein conformational changes, are being studied computationally in our group for mammalian sirtuins.

### Experimental characterization of proposed hit compounds for mechanism-based sirtuin activation

In this Section we characterize proposed sirtuin-activating compounds within the context of the mechanistic model above, evaluating their characteristics within the context of the ideal features of mechanism-based activators described above. Both thermodynamic and kinetic measurements are made in order to characterize equilibrium and steady state constants entering the model presented above.

Because such mechanism-based modulators operate according to new modes of action, as depicted in Fig. 3, the traditional approach of characterizing a compound as uncompetitive, noncompetitive or competitive with respect to substrates is not sufficient, even if the modulator has a net inhibitory effect. The models presented above provide general principles for kinetic characterization of mechanism-based enzyme activating compounds, which we apply here.

The aim of steady state characterization of hit compounds for mechanism-based activation is to estimate the parameters in equation (2) in the absence of modulator and at a saturating concentration of modulator, varying both [NAD^+^] and [NAM] in each case. This provides estimates of both the front and back face steady state parameters in Fig. 3a. By contrast, a mixed inhibition model with respect to the modulator concentration (as, e.g., done for certain mechanism-based inhibitors [17]) would not have an interpretation for the steady state constants in terms of fundamental rate constants in the sirtuin mechanism, whereas characterizing unsaturating modulator concentrations at each of multiple product (NAM) concentrations, while providing more information, would similarly estimate only apparent steady state parameters in Fig. 3b.

Along with SIRT1, SIRT3 is one of the most important sirtuins enzymes involved in the regulation of healthspan [15]; hence we chose it as a subject of study. Honokiol (HKL) has been reported as a SIRT3 activator for the MnSOD protein substrate [19]. Because SIRT3 does not share the allosteric activation site of SIRT1, we studied HKL as a potential hit compound for mechanism-based activation of SIRT3. Following dose response scans at both unsaturating NAD^+^ and peptide (Fig. 6), we carried out steady state rate measurements at multiple values of [NAD+] and [NAM] in order to estimate the parameters in model (2). The results are presented in Figs 7,8 and Table 1. Note that at very high values of [NAM], the effect of differences in K_d,NAM_ induced by the modulator becomes negligible, allowing isolation of differences in the NAD+ binding and ADP ribosylation/base exchange kinetics. Below, we compare the observed changes in the steady state parameters *v*_max_, *K_m_*_,*NAD*+_, *K*_1_, *K*_2_, *K*_3_ at 200 μM concentration (approximately saturating) to the changes conducive to activation that were delineated above.

**Figure 6.**
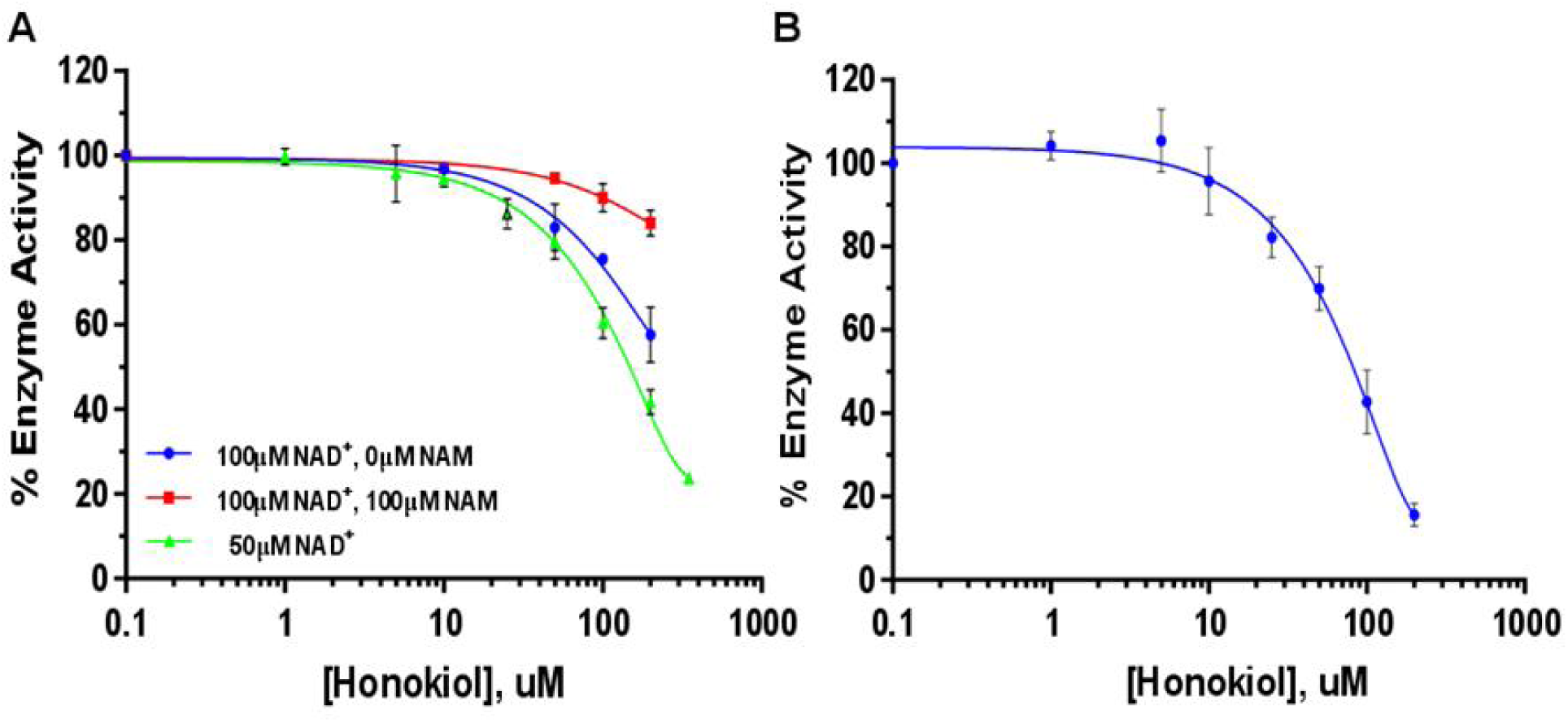
Effect of Honokiol on Sirt3 deacetylation activity using a label-free assay: dose-response. **(A)** The superposition of the un-saturating NAD^+^ and saturating MnSOD K122 peptide in the three formats of (blue) 10 mins, 100uM NAD^+^ (N=2); (red) 10 mins, 100uM NAD^+^/100uM NAM (N=2), and (green) 30 min, 50uM NAD^+^ (N=3). **(B)** 2.5 mM NAD^+^ and 6.25uM un-saturating MnSOD K122 peptide (N=5).

**Figure 7.**
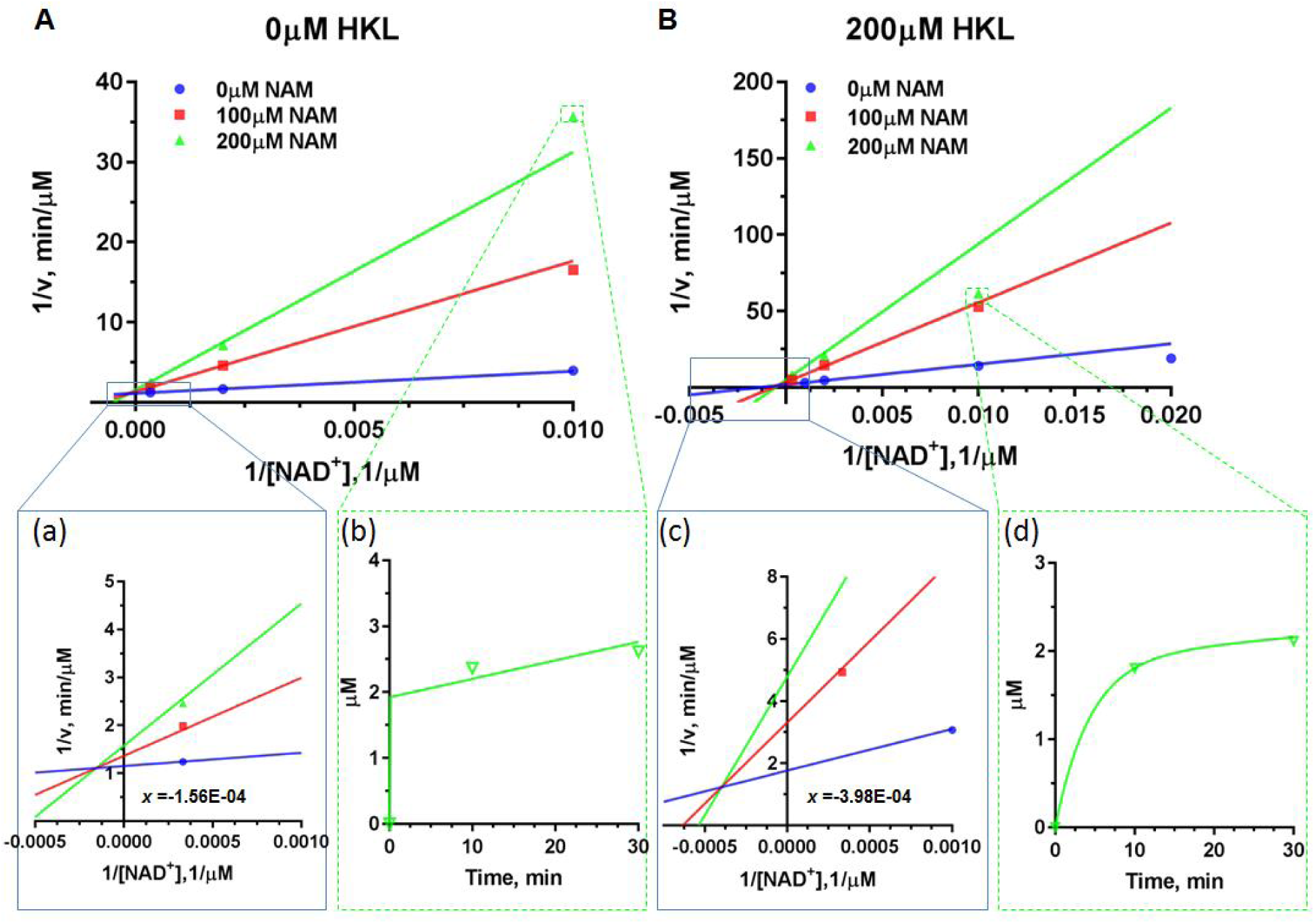
Steady-state kinetic characterization of deacetylation in the presence and absence of the mechanism-based modulator honokiol. Double reciprocal plots for deacetylation initial rate measurements of saturating substrate peptide (MnSOD K122) in the presence of different concentrations of NAM with (A) 0 μM HKL; (B) 200 μM HKL. Enlargements of the intersection points are provided as insets (a) and (c). The time series plots of mM product formed vs. time are provided as insets (b) and (d).

**Figure 8.**
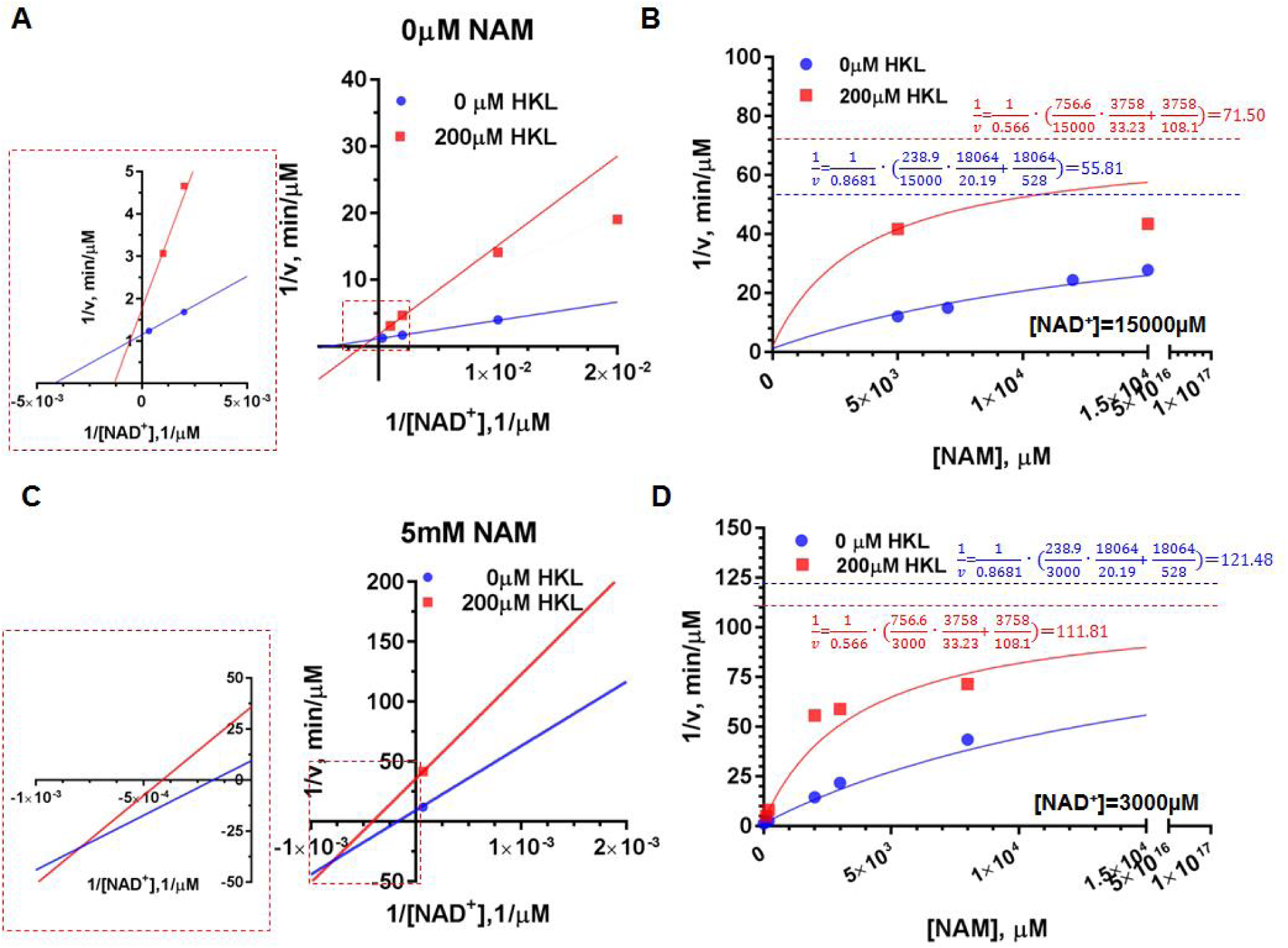
Mechanism-based modulation of Sirt3 (118-399) deacetylation of MnSOD substrate: effect of NAD^+^ and NAM. (A) Double reciprocal plots for deacetylation initial rate measurements in the presence and absence of Honokiol at [NAM]=0 μM. The enlargement of intersection point was provided as inset. **(B)** Comparison of Dixon plots at [NAD^+^]=15000 μM in the presence and absence of HKL. **(C)** Double reciprocal plots for deacetylation initial rate measurements in the presence of Honokiol at [NAM]= 200μM. The enlargement of intersection point was provided as inset. **(D)** Comparison of Dixon plots at [NAD^+^]= 3000 μM in the presence and absence of HKL. Note that the model omits the term quadratic in [NAM] in eqn (1) and the dotted lines shown in the Dixon plots are the asymptotic values to which the model-predicted rates converge in the absence of this term.

**Table 1.** Model parameter estimates from global nonlinear fitting of equation (2) for SIRT3 in the presence and absence of 200uM (saturating) HKL. [E_0_] = 1.85 μM

| | 0$\mu\text{M}$ Honokiol | 200$\mu\text{M}$ Honokiol |
| --- | --- | --- |
| Vmax | 0.8681 | 0.566 |
| K1 | 18064 | 3758 |
| Km | 238.9 | 756.6 |
| K2 | 20.19 | 33.23 |
| K3 | 528 | 108.1 |
| **Std. Error** |  |  |
| Vmax | 0.01216 | 0.05319 |
| K1 | 8279 | 1572 |
| Km | 12.92 | 138.8 |
| K2 | 1.612 | 6.886 |
| K3 | 105 | 28.98 |
| **95% CI (profile likelihood)** |  |  |
| Vmax | 0.8417 to 0.8954 | 0.4658 to 0.7421 |
| K1 | 8497 to 288496 | 1826 to 19458 |
| Km | 212.2 to 268.7 | 501.3 to 1231 |
| K2 | 16.91 to 24.25 | 20.81 to 55.33 |
| K3 | 357.7 to 946.8 | 60.92 to 243.2 |
| **Goodness of Fit** |  |  |
| Degrees of Freedom | 11 | 10 |
| R square | 0.9988 | 0.9947 |
| Absolute Sum of Squares | 0.001052 | 0.0006615 |
| Sy.x | 0.009782 | 0.008133 |

The insets in Fig. 7 show examples of the time series data collected. These insets demonstrate that deacylation of this substrate displays a two phase behavior (pre-steady state phase/steady state phase) with a relatively slow first phase. Due to this behavior, we used a two-phase rather than one-phase exponential time series fitting. Irrespective of the first phase time constant the rate in second phase is almost identical. Hence the uncertainty in steady state rates is very low. The rate is generally almost constant over the measured times in the second phase. Further discussion of time series fitting for the MnSOD substrate are provided in the Methods section.

Because of the apparent two phase time series dynamics of catalysis for MnSOD, the relative amounts of product formation in the presence vs absence of HKL depends on the choice of measurement time, with a closer correspondence at lower times. In particular, in the presence of 100uM NAM, which is on the same order of the magnitude as the physiological concentration of NAM, the amounts of product formed at 0 and 200uM HKL after 10 mins are quite close (Fig 6A). Compared to the steady state rates, the first phase rates are generally closer in magnitude.

### Measurement of binding affinities

The binding affinities of ligands to complexes in the catalytic mechanism of SIRT3/MnSOD peptide substrate were measured using microscale thermophoresis (MST; Figs. 9-14). In order to carry out binding affinity measurements on the reactive complex of enzyme, acylated peptide substrate and NAD^+^, we synthesized the catalytically inert molecule carba-NAD^+^ and used this NAD^+^ analog for those MST studies.

**Figure 9.**
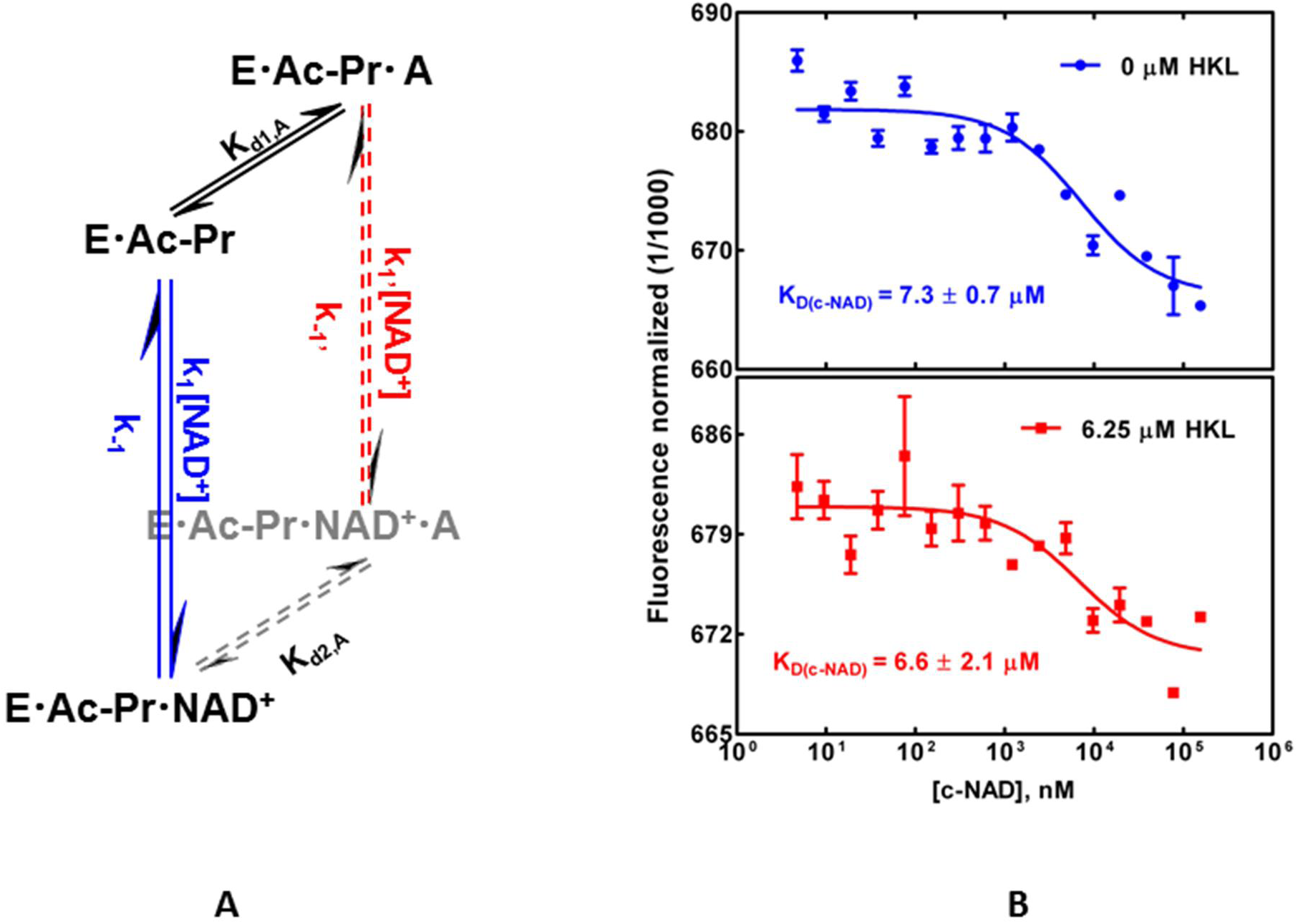
Binding affinity measurements of carba-NAD+ binding in the ternary complex: effect of mechanism-based modulator honokiol. Panel A - Pathways in the sirtuin reaction network in the presence and absence of bound modulator and inhibitor. E, enzyme; Ac-Pr, acetylated peptide substrate; NAD, nicotinamide adenine dinucleotide; A, modulator (Honokiol). Panel B - Binding of carba NAD (c-NAD) to Sirt3.Ac-MnSOD complex, in presence and absence of 6.25 micromolar Honokiol measured using MST.

In addition to measuring the effect of HKL on NAD^+^ binding affinity, we measured its effect on acylated peptide binding affinity, NAM binding affinity in the presence of acetylated peptide, deacylated peptide binding affinity, O-acetylated ADP ribose binding affinity in the presence of deacylated peptide (i.e., the product complex), and NAD binding affinity in the presence of deacylated peptide. The latter measurements in the presence of deacylated peptide were made in order to determine whether product inhibition plays any role in the observed kinetics. For binding affinity studies on complexes that appear in Fig. 3, the relevant faces of the cube are also displayed in the respective figures. Note that measuring the binding affinities for any of the 3 sides of such faces determines the fourth binding affinity. In particular, the binding affinities of HKL or carba-NAD+ were measured in the ternary complex and product complex. Cooperative binding between HKL and the relevant ligand were studied in each case. It was observed that HKL can have synergistic binding interactions with several ligands.

### Characterization of honokiol modulation of SIRT3 using kinetic and thermodynamic data

The above data indicate that HKL cobinds with the peptide substrate and cofactor of SIRT3. Under the conditions tested, the net effect of SIRT3 on MnSOD peptide substrate was inhibition, but the mechanistic model above is required to interpret the data and evaluate the compound’s viability as a hit compound for activator development.

We begin our analysis by considering the effects of HKL on SIRT3 activity in the context of traditional enzyme inhibition models. According to Fig.9, K_d,NAD+_ remains roughly unchanged for this substrate, implying that K_d1,A_ ≈ K_d2,A_. In the traditional picture, HKL binding is noncompetitive with respect to NAD^+^. According to Fig 10, the increase in HKL binding affinity in the presence of NAM suggests that K_d,NAM_ decreases for this substrate, implying that K_d3,A_ > K_d4,A_. In the traditional picture of enzyme inhibition, HKL binding is uncompetitive with respect to NAM.

**Figure 10.**
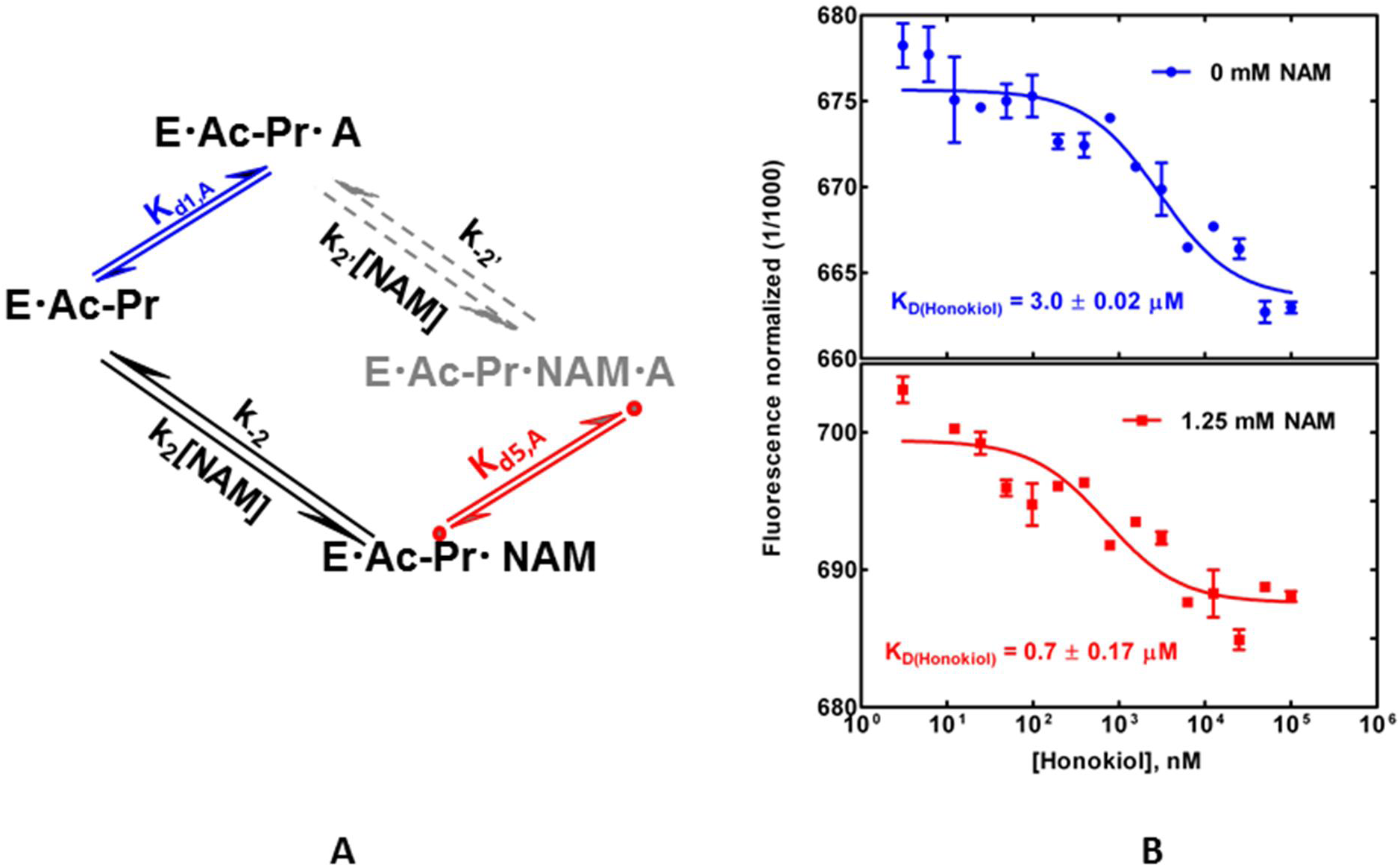
Binding affinity measurements of honokiol binding: effect of NAM. Panel A - Pathways in the sirtuin reaction network in the presence and absence of bound modulator and inhibitor. E, enzyme; Ac-Pr, acetylated peptide substrate; NAM, nicotinamide (inhibitor); A, modulator (Honokiol). Panel B - Binding of Honokiol to the Sirt3-Ac-MnSOD complex, in presence and absence of NAM, measured using MST.

However, the effect of HKL on activity can only be understood by application of the full kinetic and thermodynamic model presented above, which involves simultaneous effects of the modulator on both the forward and reverse reactions in the context of a steady-state model of the reaction. Note that in the presence of HKL, K_m,NAD+_ increases and catalytic efficiency decreases by a somewhat larger margin. The parameter estimates in Table 1 together with the full set of MST data can be interpreted within the context of the model above to shed light on the mechanism of HKL modulation and to evaluate its properties as a hit compound.

First, note that K_1_ decreases several fold in the presence of HKL, consistent with the increase in NAM binding affinity. As discussed above, a decrease in K_1_ may be inconsequential in terms of a molecule’s propensity to activate the enzyme as long as the fast k_-3_ approximation is not violated. If k_-3_ can no longer be omitted from the model, this can contribute to a decrease in catalytic efficiency (see equation 4).

Second, K_3_ decreases several fold in the presence of HKL, whereas an increase in K_3_ is desirable. However, K_1_/K_3_ remains roughly unchanged, as can be observed graphically in Fig 8b,d where the effect of K_1_ becomes diminished at high [NAM]. This suggests that the reduction in K_d,NAM_ may be playing a role in the observed reduction of K_3_(eqn 3e) Under the approximations (ii) discussed above, K_ex_ ‘may be relatively close to K_ex_. This is a favorable property of a hit compound since it suggests *K_d_*_3,*A*_ ≈ *K_d_*_2,*A*_; hit evolution could be carried out to reduce K_d3,A_. Since K_d,NAD+_ remains roughly unchanged by the modulator, if K_ex_ is also roughly unchanged, any reduction in k_2_ that affects K_m_ (see eqn 7) is associated with a similar reduction in k_-2_.

Third, *α* decreases several fold in the presence of HKL. This shifts the intersection point of the lines in Fig. 7A to the right in Fig. 7B. Under the above hypotheses that *K_d_*_2,*A*_ ≈ *K_d_*_1, *A*_ ≈ *K_d_*_3,*A*_ ≈ *K_d_*_2,*A*_, and *K_d_*_3,*A*_ >*K_d_*_4,*A*_, this is expected according to equation (12). Under approximation (ii) above, this decrease in *α* is due to the fact that K_m_ increases in the presence of HKL while K_d_ does not increase (see eqn 6). We can observe by comparing Fig 8 c,d to a,b that in the presence of NAM, the extent of inhibition by HKL diminishes, and that this effect is most pronounced at low [NAD^+^], which is the condition under which activation is therapeutically most useful. Under the above hypotheses, through further hit to lead evolution, this property might be exploited to activate the enzyme at low [NAD^+^] in the presence of physiological NAM or, going further, to allow a K_m_ decrease in the absence of product.

Next, the moderate increase in *K*_2_ (smaller than the changes in the above parameters) is consistent with equation (15) and is a direct consequence of the aforementioned changes in *K*_3_ and *α*. Finally, the observed increase in HKL binding affinity in the presence of O-acetylated ADP ribose (Fig. 11) may be consistent with the decrease in k_cat_, if product release is rate limiting for this substrate. Also, stabilization of coproduct binding may be consistent with HKL favoring a closed over an open loop conformation; for example, Ex-527, which is known to preferentially bind to a closed loop conformation, reduces coproduct dissociation rate and improves its binding affinity [17].

**Figure 11.**
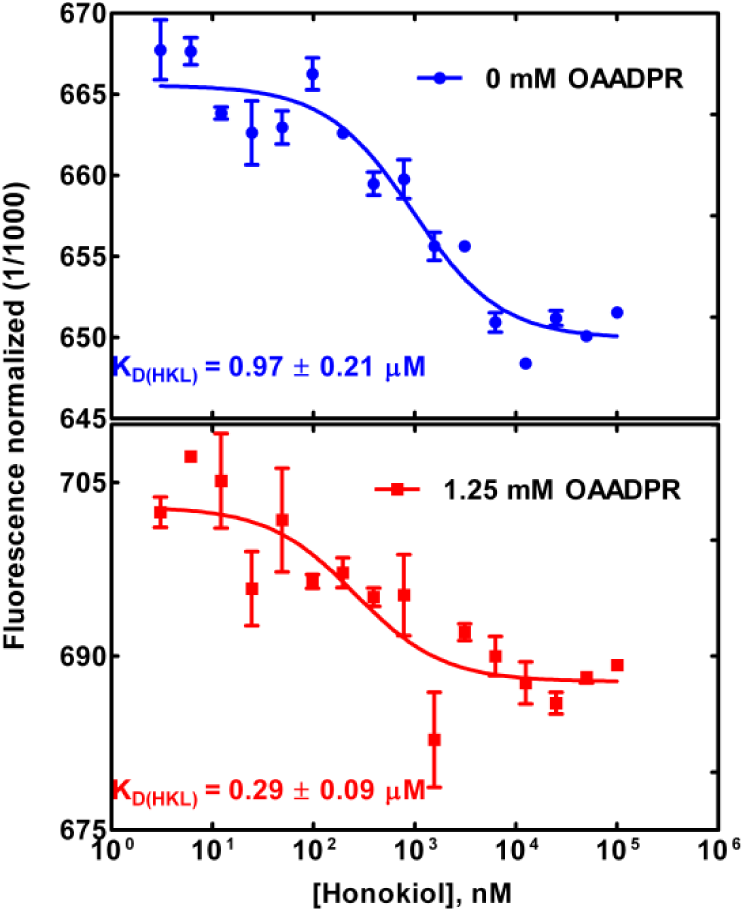
Binding affinity measurements of honokiol binding in the product complex: effect of 2’-O-acetylated ADP ribose coproduct.

**Figure 12.**
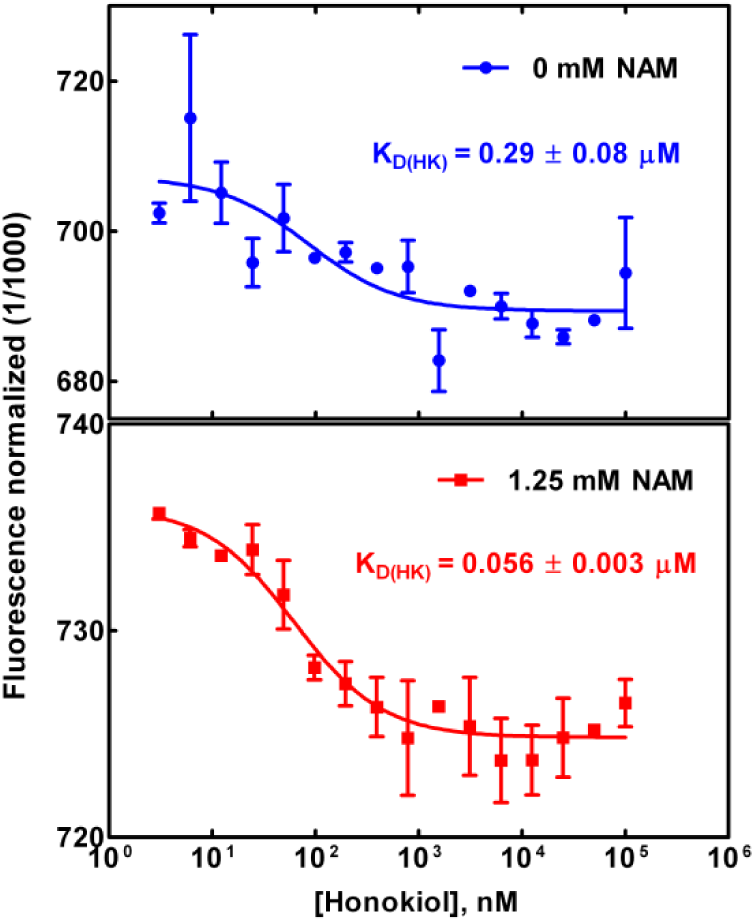
Binding affinity measurements of honokiol binding in the coproduct complex: effect of NAM.

**Figure 13.**
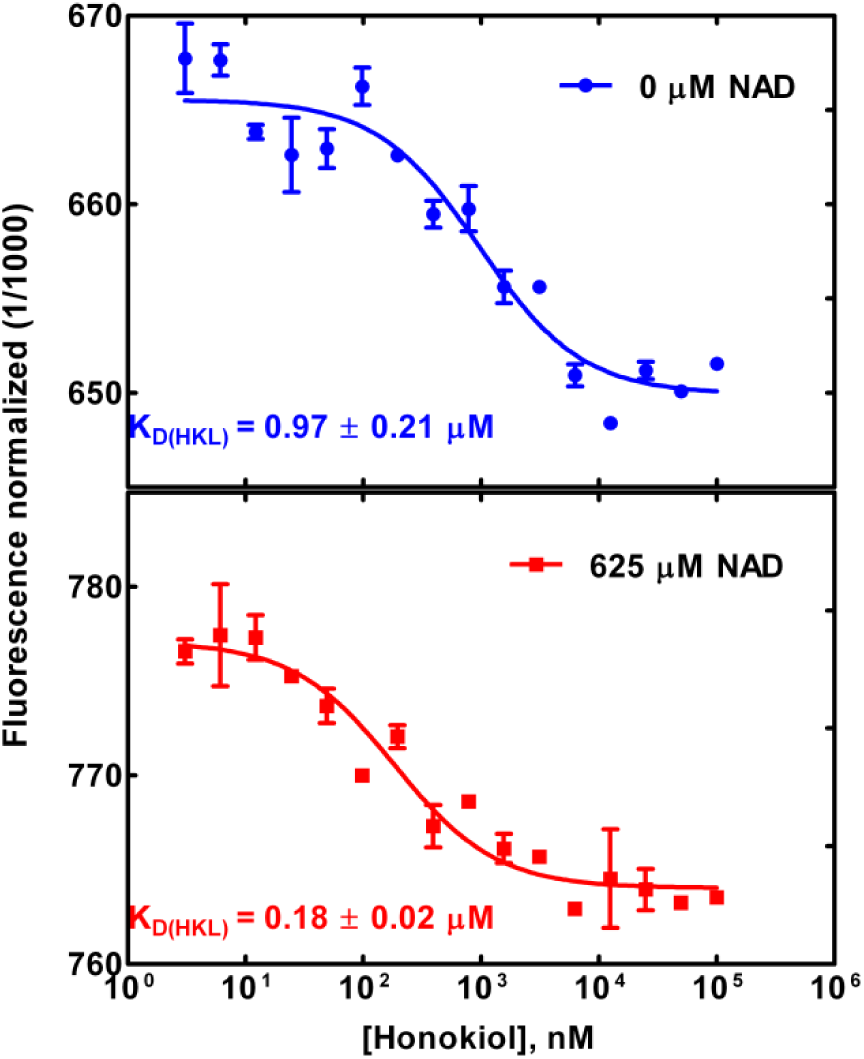
Binding affinity measurements of honokiol binding in the product complex: effect of NAD^+^.

We can also understand the properties of conventional inhibition plots with respect to modulator concentration based on the mechanistic information gleaned from our studies. Note from Fig. 8a, the intersection point of the lines with and without modulator lies to the left of the y axis and above the x axis. In the traditional mixed inhibition picture of enzyme modulation (which does not distinguish between K_m_ and K_d_), this is due to the modulator both increasing K_m_ and decreasing v_max_, but this traditional picture does not distinguish between K_m_ and K_d_, and also does not account for the relation between K_m_ and v_max_ (see eqn 4). By contrast, the equations for the lines in the presence and absence of modulator can be understood in terms of the fundamental dissociation constants and rate constants of the reaction based on our mechanistic analysis above.

Note also from Fig 8c, the intersection point of the lines with and without modulator moves closer to the x axis in the presence of NAM. For sufficiently high [NAM], the intersection point of these lines falls below the x axis and will eventually move to the right of the y axis. The fact that HKL does not increase K_d,NAD+_ gives it higher relative activity at low [NAD^+^] in presence of NAM (this is especially pronounced in the pre-steady state phase; see inset in Fig. 7d). If this could be achieved in the absence of NAM, it would signify an activator that increases catalytic efficiency while reducing v_max_.

The demonstrated ability to relate the modulated steady state kinetics of the enzyme to the manner in which a hit compound interacts with the various species in the reaction mechanism is a feature of mechanism-based activator discovery, in contrast to hit validation in traditional drug discovery, where the focus is on binding affinities [9]. By contrast, mechanism-based activation is a molecular engineering problem more akin to enzyme design.

Pre-steady state kinetics can provide additional information about the mechanism of modulation. For enzymes where product dissociation is the rate-limiting step, initial rates may not be similar to steady state rates. Hence, using initial rates in inhibition model fittings can be used for qualitative model selection (see, e.g., reference [17]), not quantitative parameter estimation. In the present work, we applied double exponential time series fits, which accounted for the pre-steady state phase, for quantitative work (see Appendix). Had we fit a single exponential, the estimated v_max_ would be similar but the estimated K_m_s would decrease, increasing the estimated catalytic efficiency. This is related to the fact that for such systems, the pre-steady state turnover rate exceeds k_cat_/K_m_* [substrate]. However, the quality of the model fit would be reduced due to the use of initial rates rather than steady state rates. A nonlinearity in the amount of product formation at low times (<= 10 min) is observed with MnSOD substrate, perhaps because higher [NAD^+^] leads to the steady state being reached more quickly.

Note that in the case of SIRT3 with MnSOD substrate, k_cat_ is approximately 2 min^-1^, which is on the same order of magnitude as the duration of the first phase in our time series experiments. Importantly, if slow product dissociation results in the significantly slower turnover rate in the second phase, then HKL may not reduce the rate of the slowest deacylation chemistry step in stage 2 of the reaction.

Having identified a hit compound for mechanism-based activation that displays at least some of the qualitative features identified above under the rapid equilibrium segments approximation, we can now use the kinetic and thermodynamic data above -- which together constitute a complete set of measurements -- to simultaneously estimate all the rate constants k_1_^’^,k_-1_^’^,k_2_^’^,k_-2_^’^,k_3_^’^,k_-3_^’^,k_4_^’^ and associated free energy changes. This will elucidate the mechanism of HKL-induced modulation in more detail than would be possible in a rapid equilibrium framework (i.e., in terms of the 7 rate constants instead of 5 *K_d_*_,*A*_’s) and will be carried out in future work, along with additional structural and biophysical characterization. The ability to identify system parameters in this manner will allow rapid characterization of mutated hit compounds during hit-to-lead evolution to identify those with a favorable balance of properties suitable for further development.

Since we did not use a full-length MnSOD protein substrate as did reference [19] in our studies, we cannot rule out the possibility of enhancement of SIRT3-catalyzed deacylation of this substrate by HKL in the absence of any NAM. Moreover, the dependence on time of the relative rates of MnSOD peptide deacylation with and without HKL warrants further study to understand the physiological effects of HKL on SIRT3 deacylation *in vivo*. These rates are more comparable at short times (pointing to the possible importance of non-steady state effects) and in the presence of NAM, which is present within intracellular compartments *in vivo*. Note that reference [19] observed an increase in SIRT3 expression levels in the presence of HKL, which may offset the effect of product inhibition especially in the limit of low [NAD^+^]. Also, we did not carry out characterization at nonsaturating peptide concentrations – as noted, analogous methods could be applied to that problem; the dose response curve in Fig. 6B shows slightly different characteristics that could be investigated further. Finally, it is also possible to carry out analogous characterization experiments at nonsaturating [HKL] in the dose response curves above, in order to obtain the apparent steady state constants defined above. The use of unsaturating modulator concentrations can allow the modulator to dissociate from the product complex, thus enabling product release. Note that the first-order rapid equilibrium segments approximations above to the steady state constants in the presence of modulator account for this and do allow for the possibility of an activity maximum at subsaturating concentrations. Dose response behavior of mechanism-based enzyme activators will be examined further in a future work.

### Substrate dependence of mechanism-based modulation

The p53-AMC substrate was studied in order to explore the substrate selectivity of modulation by HKL (Appendix). In enzyme design, the analogous problem of substrate specificity of an engineered protein is often considered. Due to the differing effects of the modulator on these two substrates, which can be identified and characterized within the present framework, substrate selectivity of mechanism-based activation can in principle be engineered.

For this substrate as well, we have verified HKL binding to catalytically active complexes (Fig. 14) and a slight cat efficiency decrease in the presence of modulator. In this case, we do not know the catalytic efficiency at higher [NAM] because we have not estimated the K_1_ parameter.

**Figure 14.**
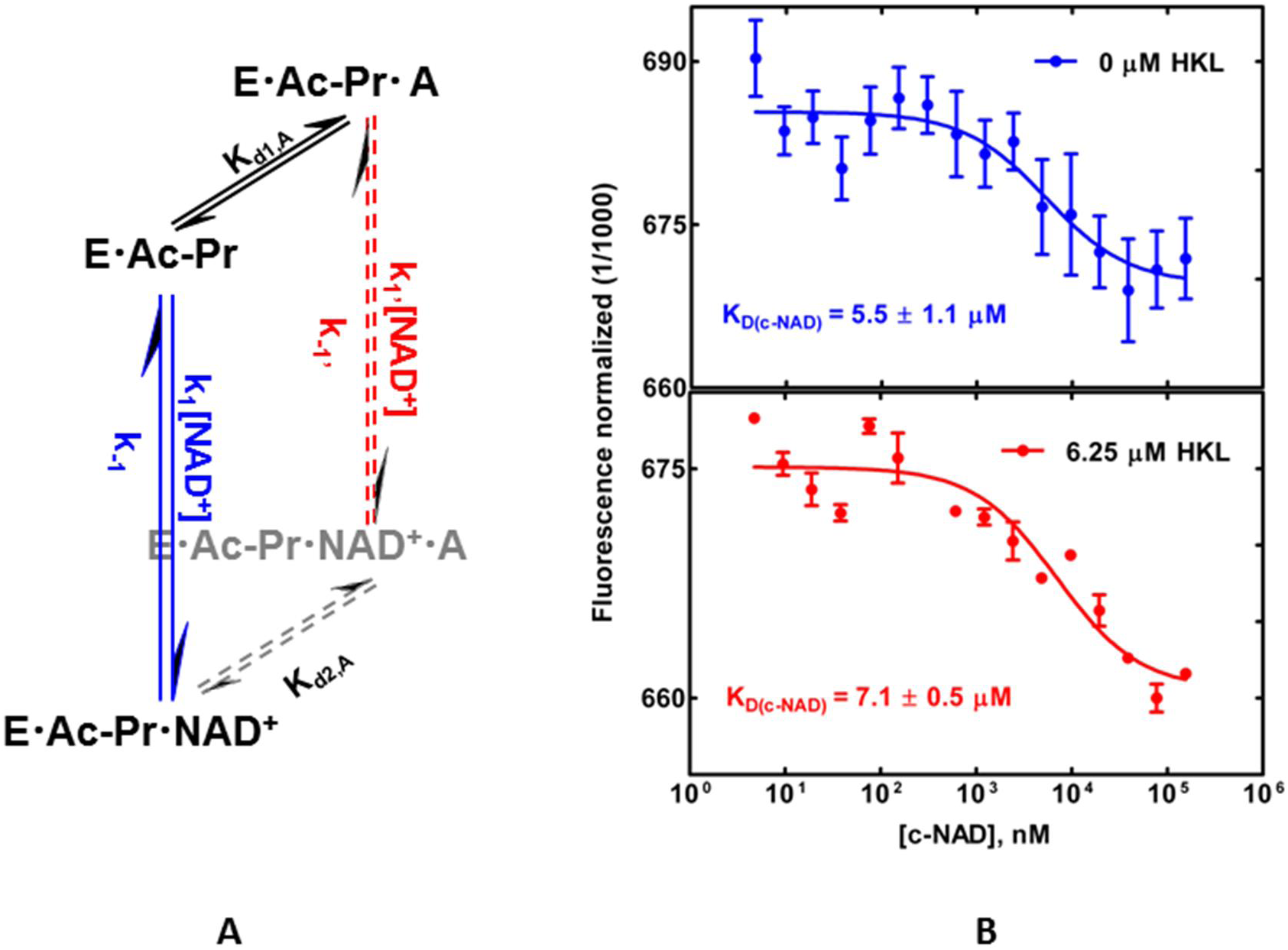
Binding affinity measurements of carba-NAD+ binding in the ternary complex of an alternate substrate: effect of honokiol. Panel A - Pathways in the sirtuin reaction network in the presence and absence of bound modulator and inhibitor. E, enzyme; Ac-Pr, acetylated peptide substrate; NAD, nicotinamide adenine dinucleotide; A, modulator (Honokiol). Panel B - Binding of carba NAD (c-NAD) to Sirt3.Ac-p53-AMC complex, in presence and absence of 6.25 micromolar Honokiol measured using MST.

In Fig. A3, which compares the initial rates of catalysis with the p53-AMC substrate in the presence and absence of saturating HKL, we observe a significant change in the y intercept (v_max_) but not K_m_ -- which would be characterized as noncompetitive inhibition in the traditional picture. However, as discussed above for the MnSOD substrate, the traditional picture is not sufficiently informative and the mixed inhibition plots with respect to NAM provide the important mechanistic information. In the absence of NAM, there is a 3-4 fold decrease in catalytic efficiency for this substrate. However, whereas *α* decreased in the presence of HKL for the MnSOD substrate, it increases for the p53-AMC substrate (Table A1). MST measurements on carba-NAD^+^ indicate a decrease in NAD^+^ binding affinity (Fig. 14). Since for MnSOD substrate, there was no change in K_d,NAD+_ and a reduction in *α*\*K_m,NAD+_ induced by HKL, the greater change in *α*\*K_m,NAD+_ induced by HKL for p53-AMC substrate observed in is consistent with the increase in K_d,NAD+_ as measured by MST.

The differences between the pre-steady state kinetics of SIRT3 for the MnSOD versus p53-AMC substrates are discussed in the Appendix.

We note that the approximations (i-iii) above, in particular (ii), may not apply to particular sirtuins and/or substrates like MnSOD peptide or p53-AMC. Their use in the analyses above were for the purpose of simplicity of illustration. The applicability of such approximations can be assessed upon full system identification – that is, simultaneous estimation of all rate constants in the model above – using a complete set of measurements. For example, without system identification, we cannot ascertain the relative magnitudes of rate constants such as those for the ADP ribosylation step (k_2_) and the rate limiting step of stage 2 (k_4_).This will be studied in a subsequent work.

In summary, HKL binds to all complexes in the sirtuin reaction mechanism, not just the product as in the case of mechanism-based sirtuin inhibitors like Ex-527. The tight binding of HKL to the coproduct does not reduce catalytic efficiency. Compounds like Ex-527 are not hits for mechanism-based activation because they do not bind to any other catalytically relevant complexes and because they reduce product dissociation rate very much, thus extinguishing the reaction under saturating conditions. Ex-527 does this for many substrates and sirtuins including AceCS2 and p53-AMC [steegborn pnas]. HKL is a hit for either substrate for reasons above.

In order to characterize the mode of action of the mechanism-based inhibitor Ex-527 [17], standard inhibition models were not sufficient and the authors had to refer to crystal structures. We have presented quantitative methods for the characterization of mechanism-based activator hits that are suitable for high-throughput studies.

### False positive testing of hit compounds

Finally, another class of reported sirtuin activators – dihydropyridines, or DHPs – were previously studied with only labeled assays [20,21] that may be prone to false positives. We studied the effects of these compounds on SIRT3 activity using the p53-AMC substrate and two assays – both a fluorescence-based and an HPLC assay. The results, presented in the Appendix, suggest that DHPs are false positive compounds and not sirtuin activators. The same labeled substrate was used by both reference [20] and us. Careful scrutiny should be applied to DHPs reported as activators, due to this finding and the autofluorescence of DHPs (see Appendix). Ideally, similar protocols for the identification of false positives will be applied to other reported activators for sirtuins other than SIRT1 studied using only labeled assays. Some previous reports of SIRT1 activation for labeled substrates did not properly identify false positives due to the lack of appropriate controls [36]. Labeled assays can be used for high throughput screening as long as appropriate controls and protocols for hit validation are regularly applied.

Application of the rigorous methods for false positive identification and characterization of mechanism-based sirtuin modulators reported herein should facilitate the elimination of false positives in highthroughput screens.

## Discussion

We have presented a model for activation of sirtuin enzymes that may be applied for the design and characterization of mechanism-based sirtuin activating compounds (MB-STACs) that can in principle activate any of the mammalian sirtuins SIRT1-7. Also, the activation model presented herein could in principle be applicable to diverse substrates, unlike previously reported allosteric activation of SIRT1 that was found to accelerate deacylation for only a small fraction of over 6000 physiologically relevant peptide substrates studied, due to the need for “substrate-assistance” in the allosteric mechanism [13]. This framework comprises a new mode of enzyme activation that is distinct from the four modes of activation previously known.

Using this modeling framework, we have shown how modulation of 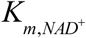 independently of 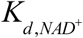 can increase the activity of sirtuins at [NAM]=0 in a manner that mimics the effects of NAD^+^ supplementation [37] but in a selective fashion. This activation also applies at nonzero [NAM], decreasing the sensitivity of the sirtuin to physiological NAM inhibition in addition to increasing its sensitivity to physiological NAD+. Such mechanism-based sirtuin activation has advantages over a) allosteric activation, which is only possible for SIRT1, is substrate-dependent, and cannot fully compensate for the reduction in NAD^+^ levels that is responsible for many aspects of health decline during organismic aging [18, 38]; b) activation of the NAD^+^ biosynthetic enzyme Nampt, which regenerates NAD^+^ from NAM and hence has nonselective effects on all enzymes that use an NAD^+^ cofactor; and c) inhibition of NAD^+^-dependent PARP enzymes, which consume NAD^+^ but are required to repair DNA damage. Moreover, it has the potential to enable isoform-specific sirtuin enzyme activation.

The example of honokiol was studied with various biochemical and biophysical methods. While further characterization is warranted, this example highlighted several important properties of hit compounds for mechanism-based activation, such as tight binding to the catalytically active complex, minimal reduction of the binding affinities of both substrates, higher relative activity in the presence of [NAM], higher relative activity under [NAD^+^] depletion conditions, and comparable pre-steady state activities in the absence of presence of modulator. The experimental evidence suggests that for MnSOD substrate (previously reported to mediate certain physiological effects of honokiol [19]), O-AADPR and NAM coproduct release are contributing to k_cat_ and catalytic efficiency reduction, respectively, with the latter being more relevant to potential therapeutic applications. Analogous methods could be directly applied to other compounds recently reported to activate sirtuins other than SIRT1, such as SIRT5 and SIRT6 [22]. The experimental characterization methods introduced were also used to identify several false positive hits reported in the literature.

The rapid equilibrium segments approximation (Appendix, equation A3) was applied in order to illustrate how a ligand that binds outside the NAD^+^ binding site can in principle increase sirtuin activity through only modulation of the relative free energies of the various species in the reaction mechanism. More detailed analysis of the mechanism of action of hit compounds for mechanism-based enzyme activation like honokiol can be achieved by complete kinetic characterization in presence/absence of the activator (e.g., by coupling base exchange with deacylation experiments). Such analyses will shed light, for example, on whether these compounds can be evolved into mechanism-based sirtuin activators that exploit the free energy profile of the sirtuin nicotinamide cleavage and base exchange reactions – and if so, how.

Structurally, binding outside of the NAD^+^ binding site (the so-called A and C pockets [25, 39]) appears to be essential for mechanism-based activation. For example, consider the binding sites of long-chain fatty acids and Ex-527[17]. We are currently exploring prospective binding sites for MB-STACs. Moreover, rational design will require analysis of the relative free energies of complexes depicted in Fig. 3. We have recently initiated computational studies [29] that assess such free energy differences for some of the front face (apo) complexes in this Figure, and further studies are in progress.

The enzyme activation theory presented herein motivates experimental workflows for the hit identification, hit-to-lead evolution, and lead optimization of mechanism-based activators. In particular, the theory enables the identification and evolution of important hits that may be inhibitors, not activators, by decomposing the observed kinetic effects of a modulator into components and identifying those molecules that display favorable values of a subset of these components as hits even if the net effect on catalytic turnover is inhibition. Compared to standard library screening for hit identification and hit-to-lead evolution, this approach allows application of multiobjective optimization techniques (through iterative mutations to functional groups) to sirtuin activator design. Hit to lead evolution could be applied, for example, to honokiol modulation of the deacylation rate of specified substrates. Such workflows would be fundamentally different from traditional drug discovery workflows and would bear more similarity to the directed evolution of enzymes. The theory presented also establishes foundations for the rational design of sirtuin-activating compounds, enabling the application of state-of-the-art computational methods to activator design in a manner analogous to computational enzyme design [40]. Once crystal structures solved, enzyme engineering methods can be used for hit to lead evolution based on above theory. For MnSOD substrate and HKL, for example, the reduction in NAM and OAADPR coproduct binding affinities could be targeted in such efforts. Finally, it raises the important question as to which enzyme families may be activatable through such a mechanism-based mode of action.

## Materials and Methods

### Chemicals and Reagents

AceCS2, MnSOD and p53 derived, acetylated peptides were synthesized from GenScript, USA. FdL peptide and DHP1 were purchased from Enzo Life Sciences (Farmingdale, NY, USA). DHP2 and carba-NAD were synthesized from KareBay Biochem, Monmouth Junction, NJ, USA and Dalton Pharma, Toronto, Ontario, Canada respectively. Honokiol was purchased from Sigma (St. Louis, MO, USA). All other chemicals, solvents used were of the highest purity commercially available and were purchased from either Sigma (St. Louis, MO, USA) or Fisher Scientific (Pittsburgh, PA, USA) or from VWR International, USA. Plasmid pEX-His-hSIRT3^102-399^ was purchased from OriGene, USA, whereas truncated version, pET21b-hSIRT3^118-399^ (also T-Sirt3) was procured from Genscript USA. Following acetylated peptides were used in this study- AceCS2- NH2-TRSG-(KAc)-VMRRLLR-COOH; p53-KKGQSTSRHK-(KAc)-LMFKTEG; MnSOD-KGELLEAI-(KAc)-RDFGSFDKF; Fluoro-peptide-(QPKK^AC^-AMC, also called p^53^-AMC), label free p^53^ peptide-QPKK^AC^.

### Solubility Measurements

Solubility of DHP-1, DHP-2, and Honokiol were measured using HPLC (Agilent 1100 series). In brief, calibration curves were established, for each compound, using concentration range covering the estimated solubility’s. The samples were prepared by adding known amount of the compounds in HDAC buffer containing a range of DMSO. The samples were allowed to equilibrate at 25°C for 48 hours before analyzing on calibrated HPLC. Over-saturated samples were prepared by adding excess compounds into the solvent mixtures of interest. The linearity was measured by R-values at least >0.99 and the estimated detection limit was around 0.002 mg/mL (2 μg/mL) based on acceptable N/S ratio.

### Carba-NAD synthesis

Carba-NAD [1((1R,2S,3R,4R)-4-((((((((2R,3S,4R,5R)-5-(6-Amino-9H-purin-6-yl)-3,4 dihydroxytetrahydrofuran-2-yl) methody)hydroxphosphoryl)oxy)oxidophosphoryl)oxy)methyl)-2,3- dihydroxycyclopentyl)-3-carbamodylpryridin-1-ium.] was synthesized according to the method described previously (Szczepankiewicz et al., 2012).

Briefly, the commercially available (1R,4S)-2-azabicyclo[2.2.1]hept-5-en-3-one (R(-)-Vince lactam), was asymmetrically dihydroxylated to provide the corresponding dihydroxy lactam after removal of the all-cis stereoisomer. Acid catalyzed methanolysis of the lactam was then carried out to give the amino ester hydrochloride. This was followed by ester reduction using lithium triethylborohydride for reduction of the ester to the alcohol. Next, the nicotinamide carba-riboside was prepared from the alcohol and 3-carbamoyl-1-(2,4-dinitrophenyl)pyridine-1-ium chloride using sodium acetate in methanol (Zincke reaction). Phosphorylation of the nicotinamide carba-riboside with POCl3 according to Lee’s procedure provided the carba-nicotinamide mononucleotide. This was coupled with adenosine 5′-monophosphomorpholidate using pyridinium to sylate and MnCl2 as a divalent cation source to create the final pyrophosphate bond.

### General method for DHP2 synthesis

The synthesis of DHP2 was carried out by a method described previously (Mai et al., 2009). Briefly, in step 1, we first synthesized the core compound, 3,5-dicarbethoxy-4-phenyl-1,4-dihydropyridines, by cyclocondensation between benzaldehyde (9.81μM), ethyl propiolate (9.92 μM) and the cyclopropylamine (14.49 μM). All three compounds were mixed together and heated at 80°C for 30 minutes in 0.5 ml glacial acetic acid. The reaction mix was allowed to cool to room temperature then mixed with 20 ml ddH2O for 60 minutes. The resulting solid product was filtered and washed with ethyl alcohol (DHP1). To obtain 3,5-dicarboxy derivatives of DHP1, we performed alkaline hydrolysis of the compound overnight at 80°C in presence of 100% ethanol. After the completion of hydrolysis, the solvent was evaporated, the residue was eluted with water (30 mL), and the resulting solution was acidified with 2N HCl. The precipitate was filtered, washed with 30 ml ddH2O three times then dried to obtain pure DHP2. The compound was then recrystallized by acetonitrile/methanol. The purity and integrity of the intermediate and final compounds were assessed by NMR.

### Expression and purification of hSirt3^102-399^

The hSirt3^102-399^ was expressed in *E*.*coli* Arctic Express (DE3) cells (Agilent Technologies). The cells were transformed and a single bacterial colony was inoculated in LB media containing 100 μg/ml ampicillin and 20 μg/ml gentamycin at 37°C as recommended by supplier. The cells were grown overnight with continuous shaking at rpm. For large scale protein purification, we inoculated 1.5% overnight grown culture into 200 ml LB medium (total 4X 200 mL) and grown at 30°C, 250 rpm for 4 hours. The protein expression was induced by adding 1 mM Isopropyl 1-thio-D-galactopyranoside (IPTG) at 15°C. After 24 hours of induction, cells were harvested by centrifugation, and the pellet was re-suspended in A1 buffer containing 50 mM NaH_2_P04, 300 mM NaCl, 10 mM imidazole, pH 8.0). The cell suspension was sonicated to lyse the cells. A clear supernatant was separated from cell debris by centrifugation at 14000 x g for 25 min at 4°C and loaded onto a 5 ml HisTrap HP column (GE Healthcare), pre-equilibrated with A1 buffer, attached to an AKTA pure FPLC system (GE Healthcare). The column was then washed with 50 ml buffer A1, followed by 50 ml buffer A2 (50 mM NaH_2_P04, 300 mM NaCl, 75 mM imidazole, pH 8.0), followed by 50 ml buffer A3 (20 mM Tris-HCl, 2M urea, pH 6.8), followed by 75 ml buffer A2. The protein was eluted with buffer B1 containing 50 mM NaH_2_P04, 300 mM NaCl, 300 mM imidazole, pH 8.0). The eluted protein fractions were pooled and dialyzed against dialysis buffer (25 mM Tris, 100 mM NaCl, 5 mM DTT, 10% glycerol, pH 7.5). The purity and the concentration of final preparation were determined by SDS-PAGE and Bradford’s reagent respectively. The purity of final protein was >85% as assessed by SDS-PAGE.

### Expression and purification of truncated hSirt3^118-399^

The T-Sirt3 plasmid was transformed, and the protein was expressed in *E. coli* Rosetta 2 (DE3) cells (Novagen) following supplier’s recommendations. For large scale purification, a single bacterial colony was inoculated in LB media with ampicillin, and grown overnight at 37°C. We inoculated 1% of overnight culture in 200 mL LB media with ampicillin and grown at 37°C, 250 rpm until the OD_600_ nm of 0.3. Then the temperature was reduced to 30°C until the OD_600_ nm to 0.6. The Sirt3 expression was induced by adding 0.3 mM Isopropyl β-D-1-thiogalactopyranoside (IPTG) at 18°C and the cell were harvested 24 hours post-induction by centrifugation. Cell pellet were re-suspended in lysis buffer containing 25 mM HEPES-NaOH, pH 7.5, 500 mM NaCl, 5 mM 2-mercaptoethanol, 5 mM imidazole, 5 mM MgCl_2_, 5 mM adenosine triphosphate, and 5% glycerol. The cell pellet was homogenized, followed by addition of freshly prepared 0.1 mM Phenylmethylsulfonyl fluoride (PMSF) and 1 mg/ml lysozyme. The homogenized cells were lysed at 4°C for 30 min followed by sonication. The cell lysate was centrifuged at 14,000 × *g* at 4°C for 25 min and the supernatant was loaded on to a 5 ml His-trap column (GE Healthcare), attached to an AKTA pure FPLC system (GE Healthcare), pre-equilibrated with lysis buffer with 20 mM imidazole, pH 7.5. The column was washed with 10 column volumes of wash buffer (lysis buffer with 75 mM imidazole, pH 7.5) and the protein was eluted using a linear gradient of wash buffer and elution buffer (lysis buffer with 500 mM imidazole, pH 7.5). The eluted protein fractions were pooled and dialyzed into lysis buffer, overnight, at 4°C. The dialyzed sample was re-loaded on to the His-trap column and subjected to a second round of purification as above. Final eluted fractions were subjected to SDS-PAGE; pure fractions were pooled together then dialyzed overnight at 4°C into dialysis buffer (25 mM Tris-HCl, 100 mM NaCl, 5 mM dithiothreitol (DTT), 10% glycerol, pH 7.5). The purity of final protein was >90% as assessed by SDS-PAGE. The concentration was determined using Bradford’s method.

### Dynamic light scattering (DLS) analysis

hSirt3^102-399^ DLS data were collected using a Dynapro Nanostar (model WDPN-08, Wyatt Technology) in collaboration with Alliance Protein Laboratories, Inc., USA) using a 1 μl quartz scattering cell at 25°C. Prior to data acquisition, the samples were centrifuged at 10,000 × g for 10 minutes. Typically 25 ten-second data accumulations were recorded and averaged to improve signal/noise. The resulting data were analyzed with the Dynamics ver 7.1.8.93 software. Mean sizes (z-average) are calculated based on the Cumulants method. Size distributions were calculated using the Dynals analysis method, with the resolution set at the default value. Weight fractions were estimated using the Rayleigh spheres model. The instrument calibration was absolute, based on units of time and distance (with distance measured by the wavelength of the light source). Since the sample was in buffer containing 10% glycerol, the viscosity and refractive index of the buffer were assumed to be equivalent to those for 10% glycerol within the 1-2% precision of this technique.

### Size exclusion chromatography (SEC) analysis

For SEC analysis, a superdex-200 column (1 × 30 cm, GE Healthcare, Alliance Protein Laboratories, Inc., USA) was equilibrated with 0.1 M Na phosphate, 0.2 M arginine, pH 6.8 at a flow rate of 0.5 ml/min or with 0.1 M Na phosphate, 0.5 M NaCl, pH 7.2 at a flow rate of 0.2 ml/min. Total 400 μg T-Sirt3 protein was loaded onto the column. The protein was eluted with the same buffer. The protein concentration in each fraction was monitored at 280 nm.

### hSirt3-ligand binding assay by Microscale Thermophoresis (MST)

The hSirt3 protein was conjugated with Alexa 647 fluorophore (ThermoFisher Scientific) by NHS ester chemistry using manufacturer’s recommendations. Briefly, the conjugation reaction was performed in 20 mM HEPES, 200 mM NaCl, 0.5 mM TCEP at pH 7.5 buffer. In order to label at-least one lysine per protein molecule, a 2:1 molar excess of reactive dye was used over protein. Unconjugated dye was removed using a size exclusion chromatography and the labeled protein (Sirt3 NT647) was buffer exchanged into 50 mM Tris-HCl pH 8.0, 137 mM NaCl, 2.7 mM KCl, 1 mM MgCl2, 5% DMSO, 0.05% Pluronic F-127. A final concentration of 2 nM hSirt3 NT647 was titrated with varying concentrations of the modulator. The thermophoresis was measured (excitation wavelength 650 nm, emission wavelength 670 nm, LED-power 15%, laser-power 80%) using a Monolith NT. 115 Pico (NanoTemper Technologies) at 25°C in the absence and presence of various concentrations of NAD+, acetylated and de-acetylated peptide (as described in respective figure legends). Dissociation constants (K_D_ were determined with GraFit7 (Erithacus Software) by nonlinear fitting using a 1:1 binding model.

### Effect of Honokiol on hSirt3^102-399^ deacetylation activity

Reactions were performed in triplicate which included either 2.5 mM NAD^+^ and 6.25 μM MnSOD derived synthetic peptide or 50 μM NAD^+^ and 600 μM peptide substrate in presence of different concentrations of Honokiol (Catalogue # H4914, Sigma), ranging from 0-200 μM, in a buffer containing 50mM TRIS-HCl, 137mM NaCl, 2.7mM KCl, and 1mM MgCl2, pH 8.0 and 5% DMSO. The reactions were started by addition of 5U (**1.85 μM**) of hSirt3^102-399^ and incubated at 37°C for 30 minutes. The reactions were terminated by adding 2% TFA. The peptide product and substrate were resolved using HPLC.

### General HPLC method for peptide product separation

#### Separation of MnSOD peptide

We used an Agilent 1260 infinity HPLC system and a ZORBAX C18 (4.6x250 mm) column were used throughout the study. Deacetylated product was formed from the enzymatic reactions were separated using gradient comprising 10% acetonitrile in water with 0.005% trifluoroacetic acid (solvent A) and acetonitrile containing 0.02% TFA (solvent B) using a constant flow rate of 1 ml/min. After injection of the sample, HPLC was run isocratically in solvent A for 1 min followed by a linear gradient of 0-51% solvent B over 20 minutes.

#### Separation of other peptides

A Beckman System Gold HPLC and a ZORBAX C18 (4.6x250 mm) column were used for other peptides. Deacetylated product and substrate were separated using gradient system comprising 0.05% aqueous trifluoroacetic acid (solvent A) and acetonitrile containing 0.02% trifluoroacetic acid (solvent B) using a constant flow rate of 1ml/min. Upon injection of the sample the HPLC was run isocratically in solvent A for 1 min followed by a linear gradient of 0-51 %B over 20 min.

In both the cases, after completion of the gradient, column was washed with 100% solvent B and then equilibrated with 100% solvent A. The detector was set at 214 nm to detect the deacetylated peptide product, and the substrate. The amount of product produced was derived from the percent of product generated during the reaction. The percent of product produced was calculated by dividing the product peak area over the total area. We used GraphPad Prism software (GraphPad Software, Inc, CA) to fit the data.

### Effect of DHP1 on hSirt3^102-399^ deacetylation activity

To assess the effect of DHP1on hSirt3^102-399^ deacetylase activity, we incubated 5U hSirt3 with 500 μM NAD+, 250 μM fluorolabeled peptide (FdL2 peptide, Enzo Life Sciences Farmingdale, NY, USA) with varying concentrations of DHP1 (0-100 μM) in a buffer containing 50 mM Tris/Cl, pH 8.0, 137 mM NaCl, 2.7 mM KCl, 1 mM MgCl_2_, and 1 mg/mL BSA. The reactions were carried out for 60 minutes at 37°C and terminated by adding 1X Developer (Enzo Life Sciences). The fluorescence produced was measured (Excitation/Emission: 355/455 nm) using TECAN plate reader (TECAN Infinite M200 PRO, Switzerland, Tecan Group Ltd). For measuring the hSirt3^102-399^ deacetylase activity in presence of both NAM (50 μM) and DHP1 (0-50 μM), the reactions were carried out as described above but the FdL2 peptide concentration was changed to 100 μM.

### hSirt3^102-399^ kinetic analysis using Fluorolabeled Peptide

We determined the steady state kinetic parameters of hSirt3^102-399^ deacetylation using fluorolabeled FdL2 peptide. The enzymatic reactions were carried out similar to as described above. For initial velocity determination, we performed the reaction at different NAD^+^, and 100 μM FdL2 peptide. We terminated the reactions at specified time by adding the 1X developer. After the completion of the reaction, the fluorescence intensity was measured on TECAN microplate reader. The raw data were fitted to the model equations equation using GraphPad Prism (GraphPad Software, Inc, CA) to obtain the kinetic constants.

### Effect of DHP1 and 2 on hSirt3^102-399^ deacetylation activity

Sirt3 enzyme reactions were performed to assess the effect of DHP1 and 2 using p53 derived peptide. We incubated different concentrations of DHP2 with either 3 mM NAD^+^ and 10 μM p53 derived unlabeled peptide or 3 μM NAD^+^ and 250 μM peptide substrates in presence of different concentrations of DHP2, ranging from 0-400 μM. The reaction buffer contained 50 mM TRIS-HCl, 137 mM NaCl, 2.7 mM KCl, and 1 mM MgCl2, pH 8.0. The reactions were started by addition of 5U hSirt3^102-399^ and incubated at 37°C for 30 minutes. The reactions were terminated by adding 2% TFA final then were resolved on C18 column as described above. For reaction with DHP1, 50 μM of DHP1 was used.

## Appendix

### Time series fitting

For the MnSOD substrate, a single exponential model fit poorly at times between 0 and 40 mins. The time series data displayed a two phase behavior, one phase between 0 and up to 10 mins, the latter being the measurement time of the first data point. For this system, a single exponential fit would average properties of the first and second phase s, which would not provide an accurate steady state rate estimate. Ultimately the sampling frequency is too low for more accuracy than a double exponential fit and for accurate estimation of the time constant of convergence to the steady state. Example double exponential fittings are shown in the insets of Fig. 7.

In this regard, further analysis is needed to determine whether there is dynamic pre-steady state activation in the presence of HKL, for example through carrying out non-steady state experiments at high [E]_0_ and low [NAD^+^].

Also, in order to assess the accuracy of steady state rate calculations for this system, we tested for product inhibition by adding exogenous O-acetylated ADP ribose to the system and evaluating the effect on the time series curve. The results are presented in Fig. A2.

**Figure A1.**
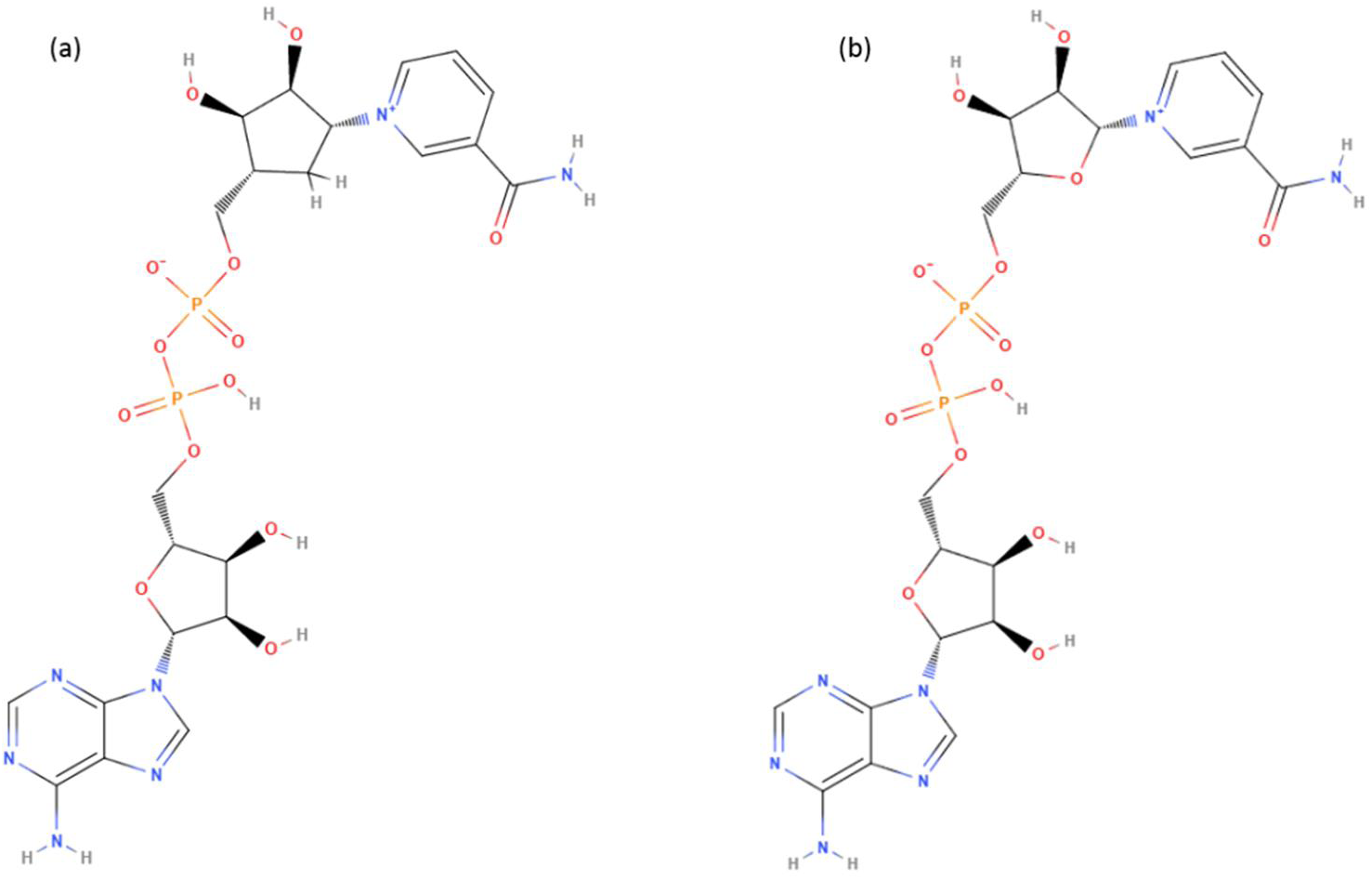
Nicotinamide adenine dinucleotide chemical structures. (A) c-NAD^+^; (B) NAD^+^

**Figure A2.**
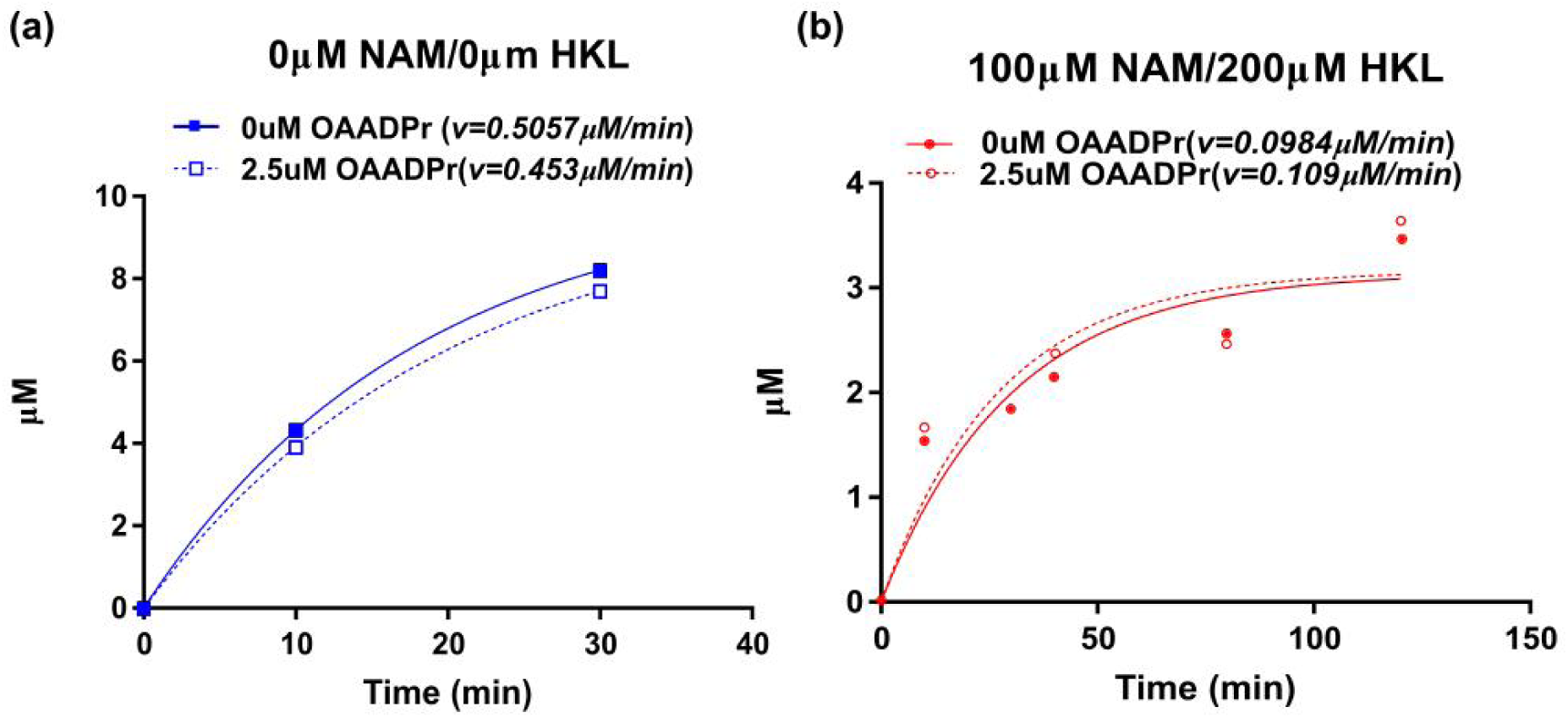
Investigation of OAADPr product inhibition. The deaceylation reactions were performed at [NAD^+^]=100μM, [MnSOD K122]=600μM in the presence of 2.5 μM OAADPr **(A)** 0μM NAM and 0μM HKL **(B)** 100μM NAM and 200μM HKL. The product formation was monitored by HPLC at desired time points (0, 10, 30, 40, 80, 120min). The time series were fitted in single exponential function in Prism.

Comparing the effects of saturating HKL and saturating Ex-527 [17] on SIRT3 activity, due to the substantial reduction in coproduct dissociation rate (steady state k_cat_ close to 0), saturating Ex-527 effectively constitutes “single hit” conditions wherein each enzyme can only turnover products once. This is not the case for saturating HKL, where activity remains.

Following system identification of all rate parameters in the steady state model, the rate constant for the limiting chemistry step in stage 1 can be identified based on non-steady state experiments and the analytical solution for the time series dynamics. The time series data and model fittings suggest that at low [NAD^+^], the pre-steady state turnover rates are faster than the turnover rates at steady state and implied by k_cat_/K_m_. This results in the initial deacetylation rates under these conditions being much closer in the absence vs presence of HKL compared to the estimated steady-state rates. Note that k_cat_/K_m_ is only applicable under steady state conditions and not at pre-steady conditions such as those at small times or very low [NAD^+^]. Although we cannot conclusively state the causes of the closer rates of MnSOD deacetylation in the pre-steady state phase without further analysis, the fact that SIRT3/MnSOD has a fast pre-steady state phase would mean product release rate improvement is a goal of hit to lead evolution.

Note that the pre-steady state phase is not detectable from time series data for the p53-AMC substrate. It is possible that product release is not rate limiting for this substrate. In that case we would expect the reduction in v_max_ to be caused by reduction in the rate constant for the slowest chemistry step in stage 2; but as noted in the text, evidence was to the contrary for MnSOD substrate. Alternatively, if product release is rate limiting, the latter chemistry step rate may be closer in magnitude to that of product release than in the case of MnSOD.

### Substrate dependence of HKL modulation of SIRT3

**Table A1.**
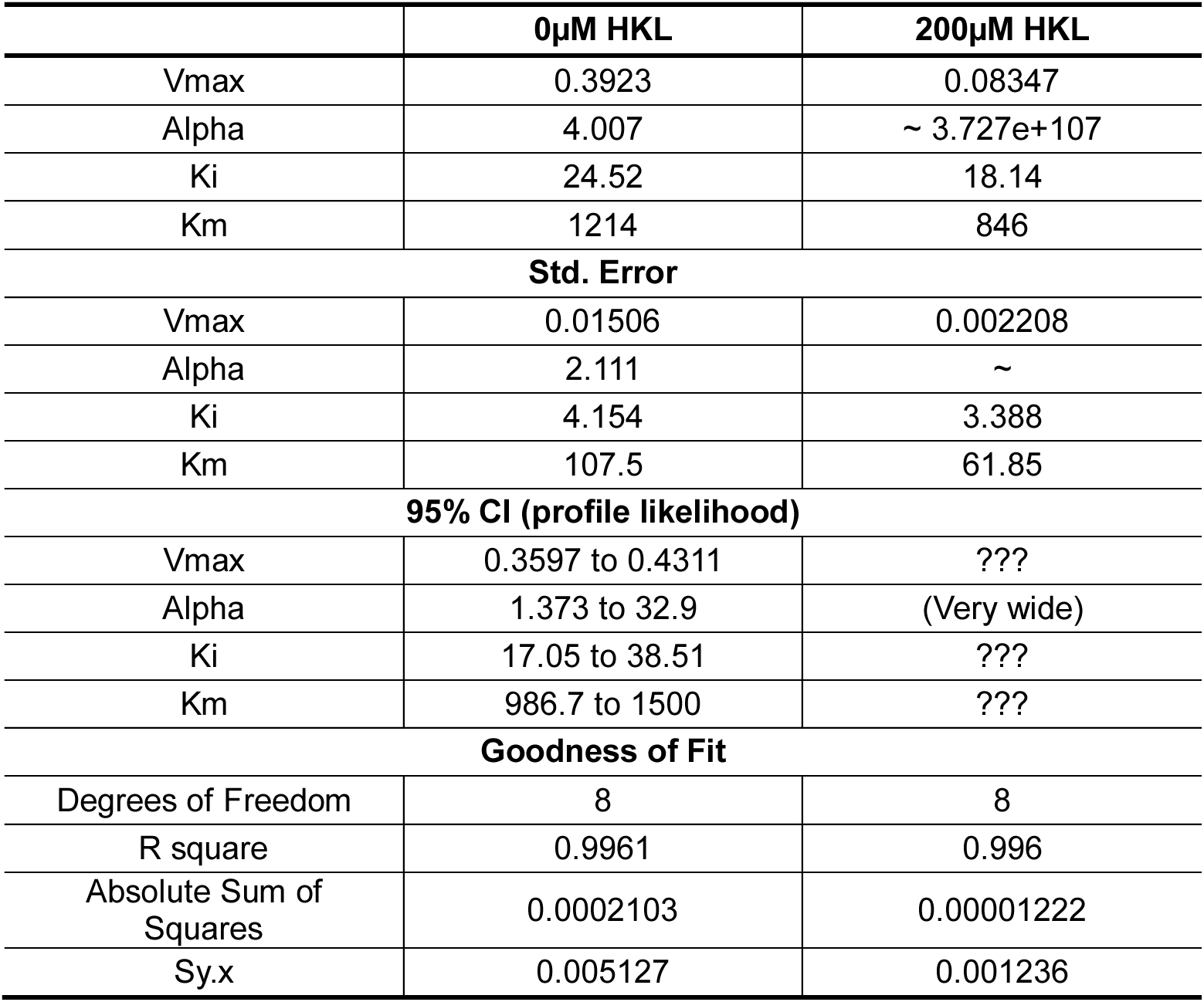
Model parameter estimates from global nonlinear fitting of mixed inhibition for SIRT3-FdL in the presence and absence of HKL. [E_0_] = 3.05 μM

**Figure A3.**
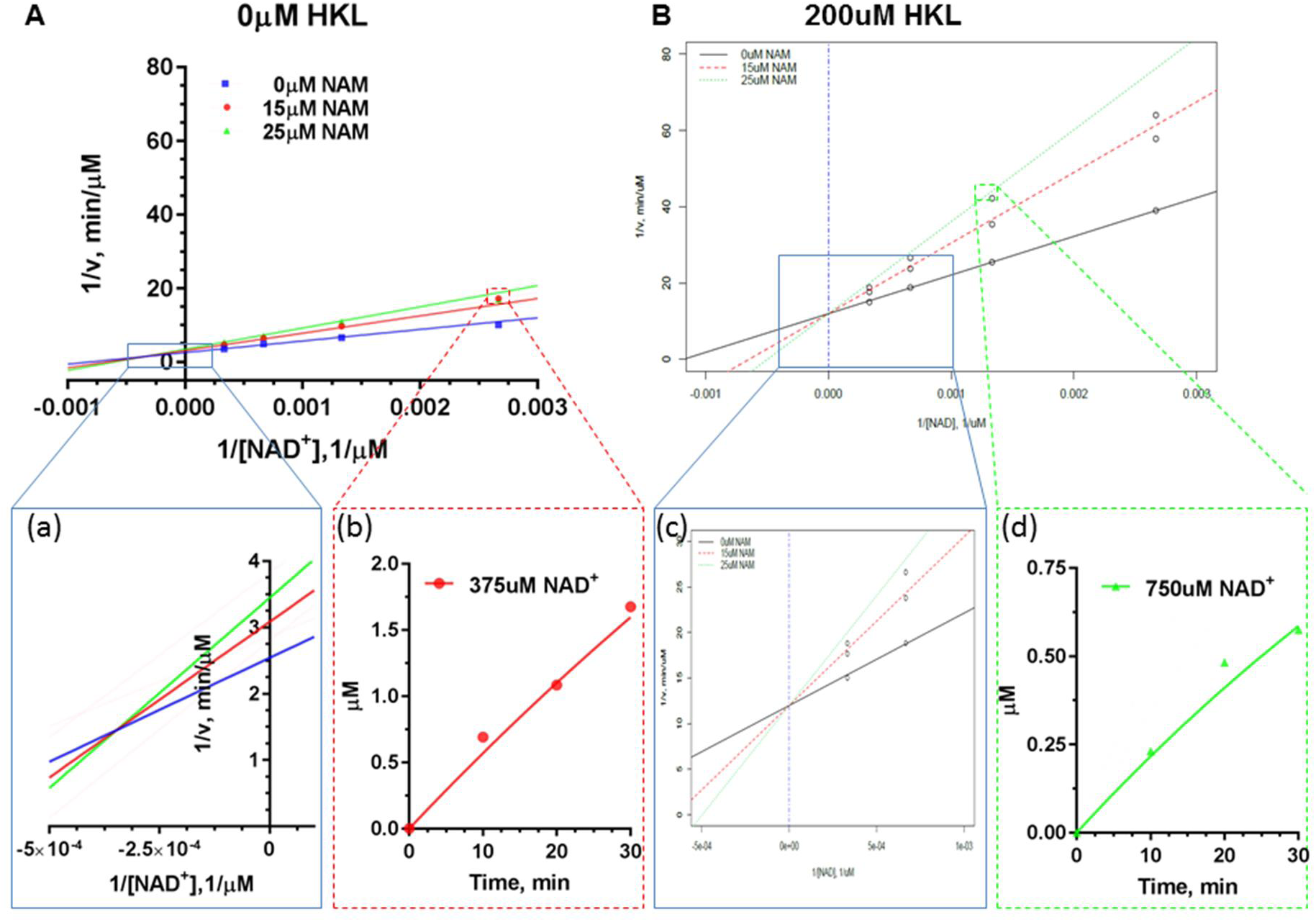
Double reciprocal plots for deacylation initial rate measurements of saturating substrate peptide (FdL) in the presence of different concentrations of NAM with (A) 0 μM HKL; (B) 200 μM HKL.The enlargement of the intersection points were provided as insets (a) and (c). The time series plot of mM product formed vs. time were provided as insets (b) and (d).

**Figure A4.**
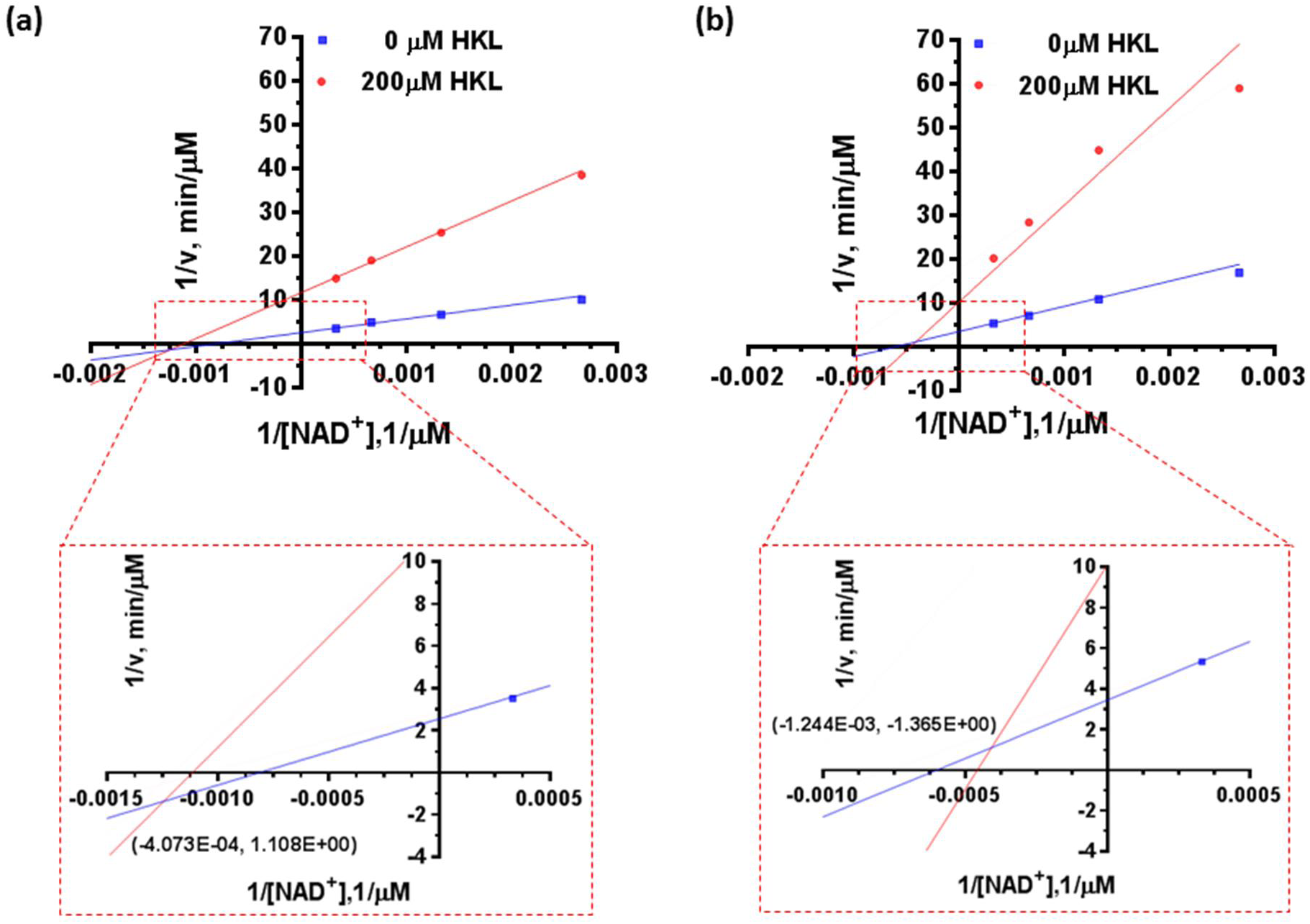
Double reciprocal plots for deacylation initial rate measurements of saturating substrate peptide (FdL) in the absence and presence of Honokiol with **(A)** 0 uM NAM; **(B)** 25uM NAM. The zoom in of insect point with x,y values were provided as insets.

### False positive testing of labeled assays

**Figure A5.**
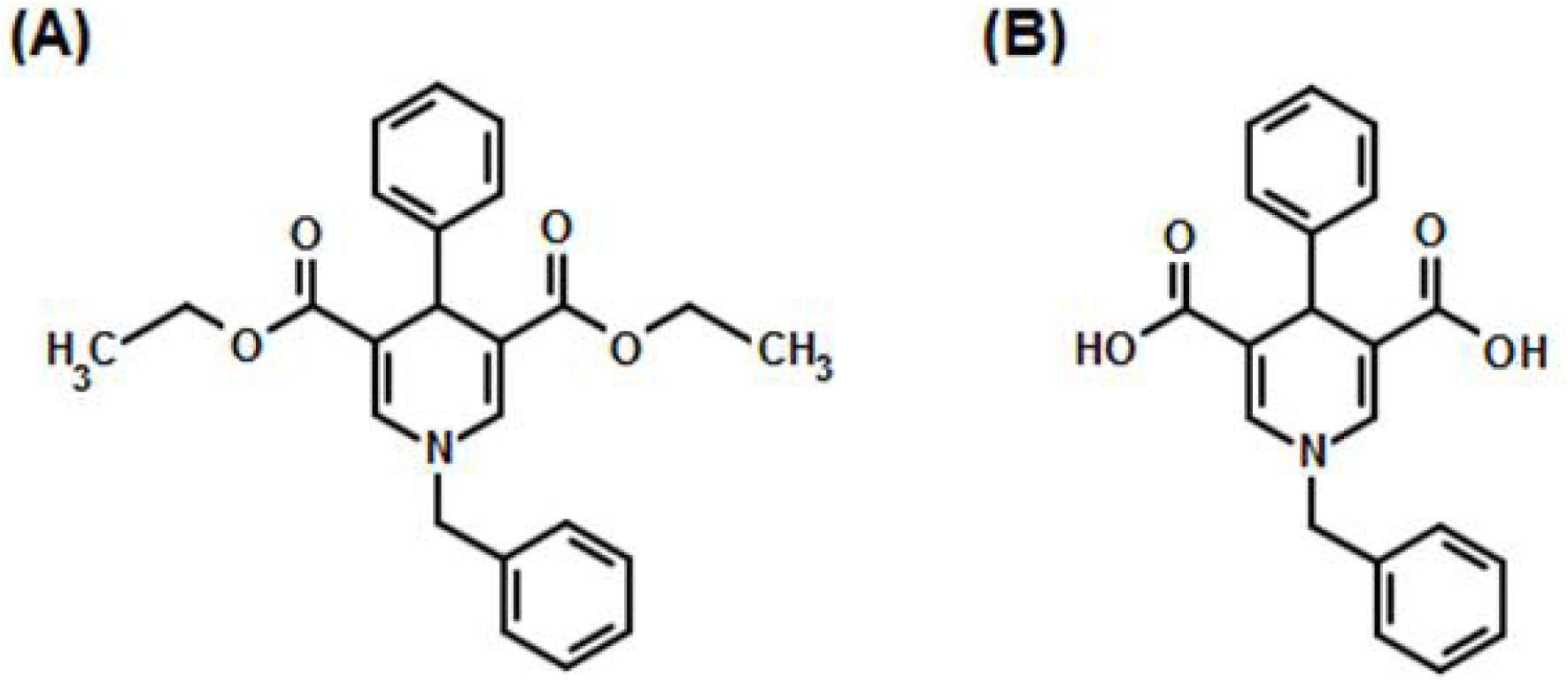
Chemical structures of the sirtuin test compounds. **(A)** N-Benzyl-3, 5-dicar-bethoxy-4-phenyl-1, 4-dihydropyridine (DHP-1). **(B)** N-Benzyl-3, 5-dicarboxy-4-phenyl-1, 4-di-hydropyridine (DHP-2).

DHP-1 (Fig A5A) can be dissolved in reaction buffer at 50uM, but only in the form of a metastable solution. Measurement of DHP-1’s solubility using the above protocol revealed that it is thermodynamically insoluble. By mutating the ester groups in DHP-1 to carboxylic acid groups, we obtain the mutated compound DHP-2 (Fig. A5B). In contrast to DHP-1, DHP-2 is thermodynamically soluble (Table A2):

**Table A2.** Solubility of DHP-2 in different % DMSO/HDAC solution.

| Compound | Solvent | AUC | Solubility (mg/mL) |
| --- | --- | --- | --- |
| 1-benzyl-4-phenyl-1,4-dihydropyridine-3,5-dicarboxylic acid<br>(DHP2) | 100% HDAC | 21397 | 2.303 |
|  | 2% DMSO | 16212 | 1.745 |
|  | 5% DMSO | 19867 | 2.138 |
|  | 10% DMSO | 20392 | 2.195 |

The solubility of Honokiol was also assessed with this protocol (Table A3):

**Table A3.** Solubility of Honokiol in different % DMSO/HDAC solution.

| Compound | Solvent | AUC | Solubility (mg/mL) |
| --- | --- | --- | --- |
| Honokiol | 100% HDAC | n.d. | n.d. |
|  | 1% DMSO | n.d. | n.d. |
|  | 2% DMSO | 260.8 | 0.041 |
|  | 5% DMSO | 766.2 | 0.120 |

Autofluorescence contributes to what is apparently false-positive activation of Sirt3 by the DHP-1 and DHP-2 compounds. A recent study [20,21] suggested that DHP-1 and 2 were a potent Sirt3 activator using FdL drug discovery kit. However, our results showed that DHP-1 and 2 emit light at wavelengths of 440- 460nm, same as the fluorophore (AMC) used in the FdL drug discovery kit (Fig. A8). The strong autofluorescence signal interferes with the readout from the real deacetylation reaction. Standard curves in the presence of different concentrations of DHP-1 (not shown) revealed that (1) the slope changes with varying concentration of [DHP-1]; (2) poor linear correlation was observed at higher [DHP-1]s. Those all indicated that the subtracting baseline fluorescence background did not clear out the autofluorescence effect. Similar effects were observed for DHP-2, where reference [20,21] reported activation under the conditions tested here (presumably using a single standard curve), whereas we observed apparent inhibition using the FdL assay at these and higher concentrations using a concentration-dependent standard curve. Fig. A7 shows the divergence between dose response curves collected for DHP-2 using a label-free HPLC assay vs the labeled FdL assay. The HPLC assay showed no significant activity modulation at any of the tested concentrations. This divergence was not observed for HKL, which does not display autofluorescence at the detection wavelength of the FdL assay (Fig. A6).

**Figure A6.**
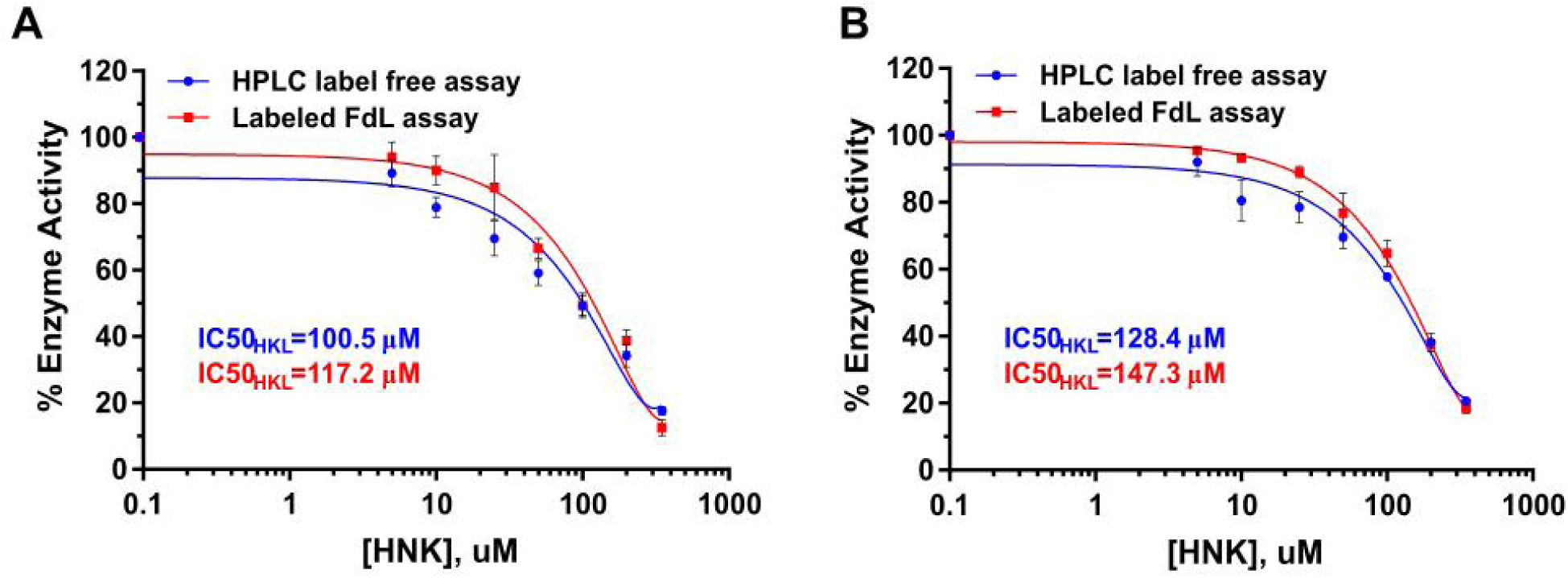
Comparison of dose response of HKL for FdL peptide using label-free HPLC assay and labeled FdL assay. **(A)** Saturating FdL peptide (250uM) and un-saturating NAD^+^(25uM) (N=2); **(B)** high NAD^+^ (3000uM) and un-saturating FdL peptide (30uM) (N=2) in the presence of different concentrations of HKL range from 0-350uM using label-free HPLC (red) and labeled FdL assay (blue).

**Figure A7.**
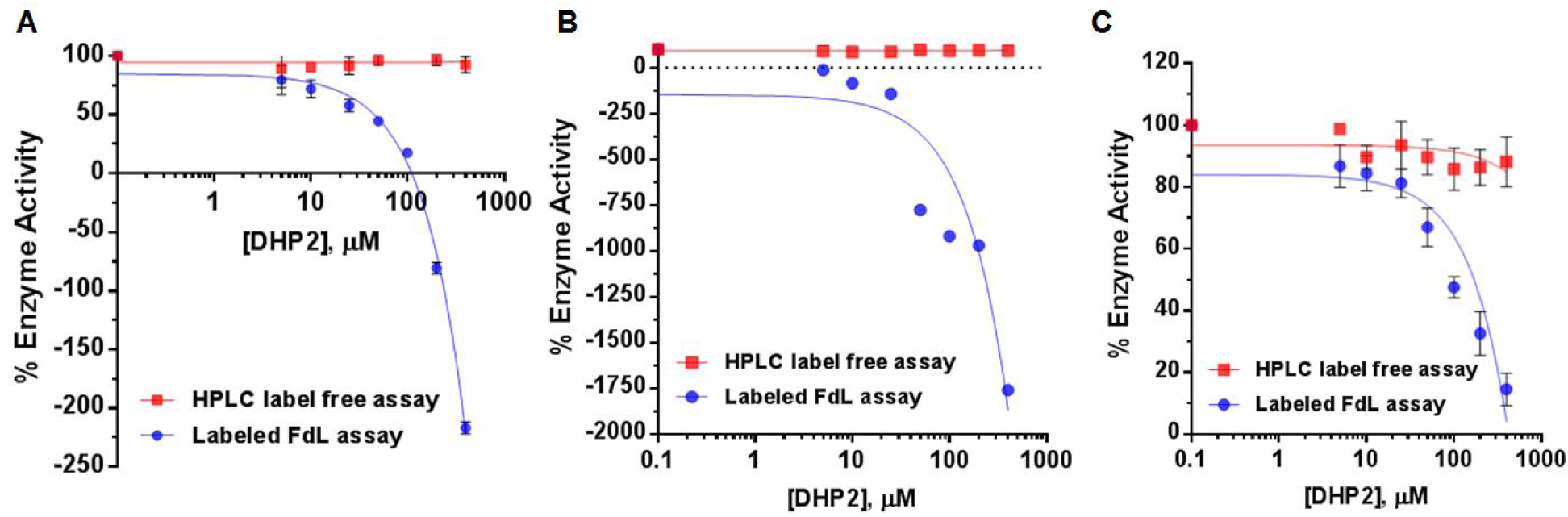
Comparison of dose response of DHP2 for FdL peptide using label-free HPLC assay and labeled FdL assay. **(A)** Saturating FdL peptide (250uM) and un-saturating NAD^+^ (10uM) (N=2); **(B)** high NAD^+^ (3000uM) and un-saturating FdL peptide (3uM) (N=2) in the presence of different concentrations of DHP2 range from 0-400uM using label-free HPLC (red) and labeled FdL assay (blue).

**Figure A8.**
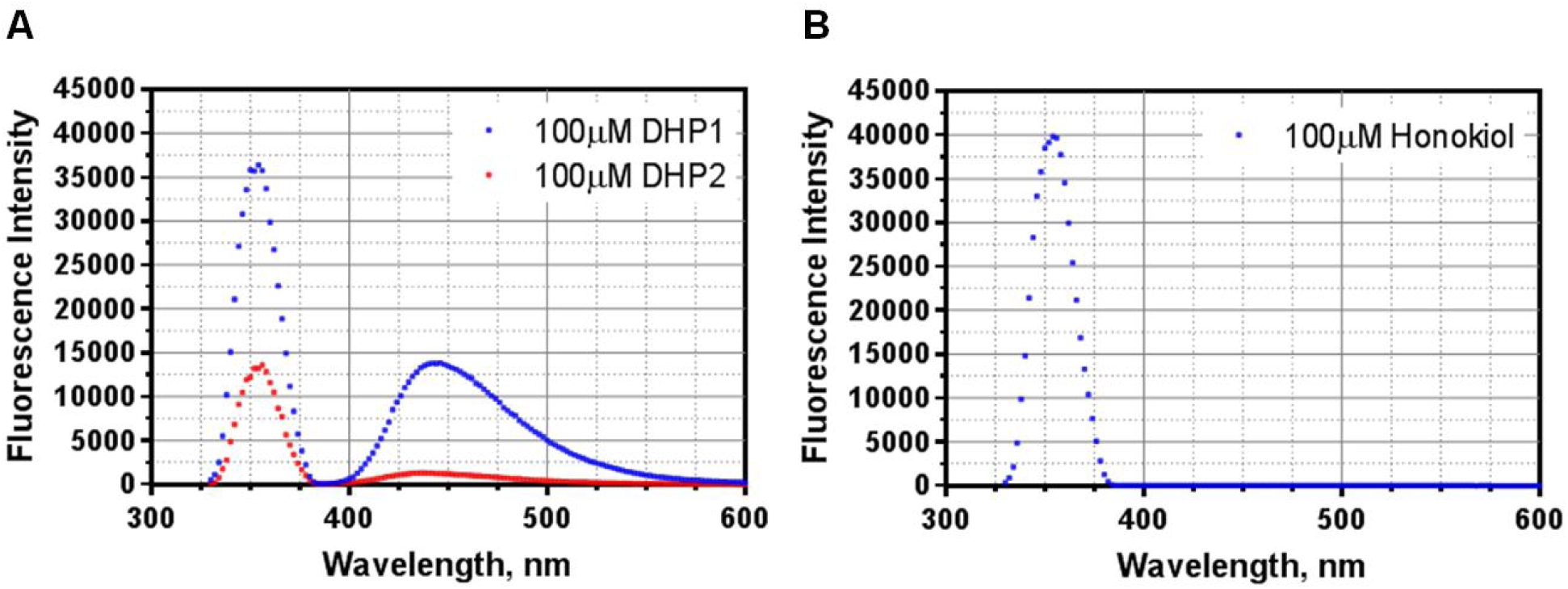
Auto-fluorescence scan for **(A)** DHPs and **(B)** HKL were performed on a multifunctional microplate reader (TECAN Infinite M200 PRO, Switzerland, Tecan Group Ltd.) with setting of Ex=355nm, Em=330-600nm, and Gain = 45. The 100uM of modulator sample solutions were prepared in either DMSO (DHP1/HKL) or HDAC (DHP2) based on their solubility (Table 3 and 4).

Note that the Sirtainty assay used in [21] also displays autofluorescence in the target wavelength range.

Expressions for c_ij_’s in equation (1):

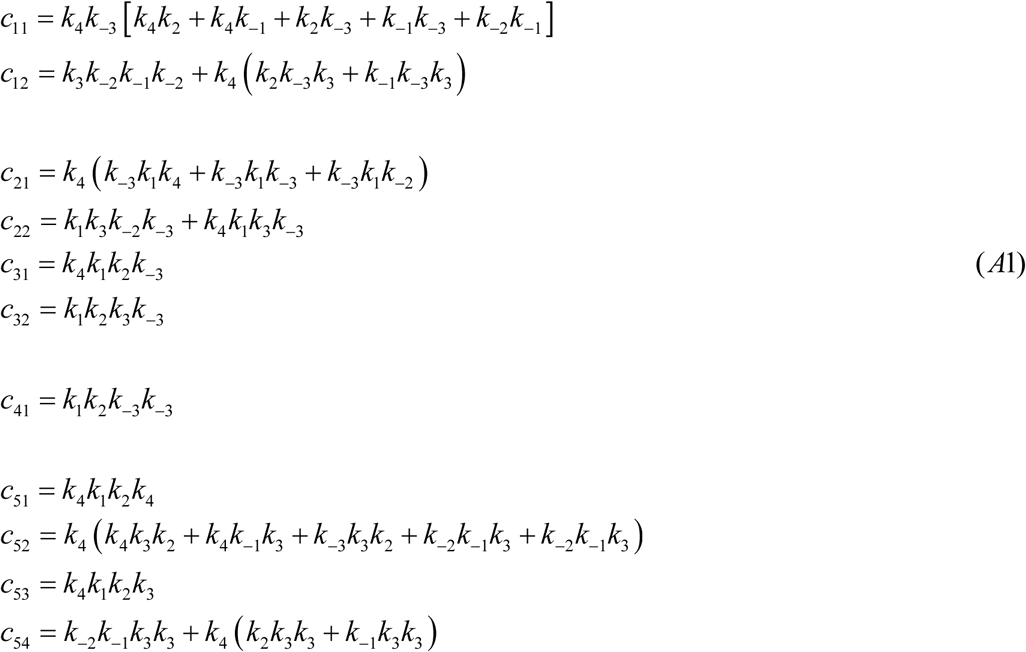

With these, the expressions for the steady state constants in equation (3) follow from

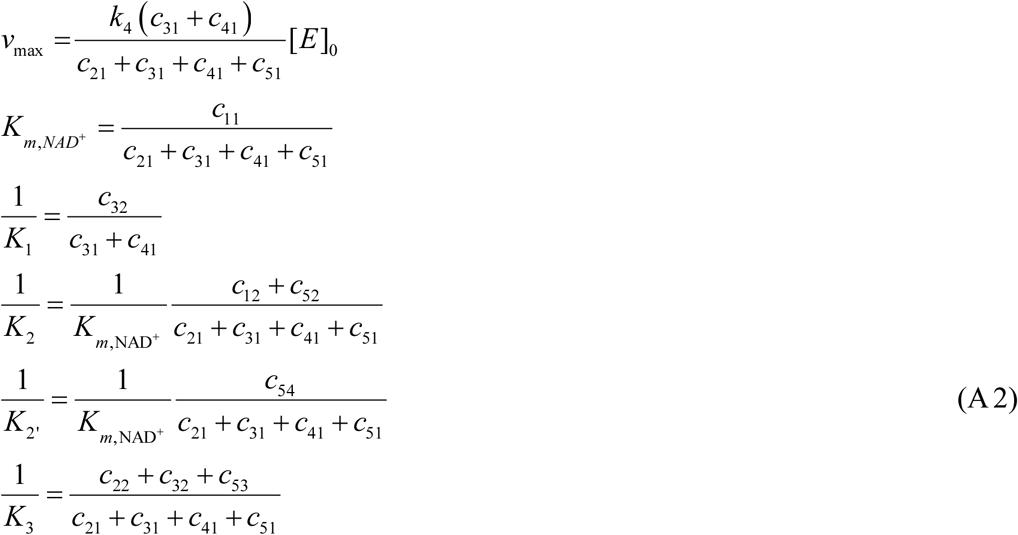

The rapid equilibrium segments expressions for the steady-state concentrations of the species in the sirtuin reaction mechanism depicted in Fig. 3 are:

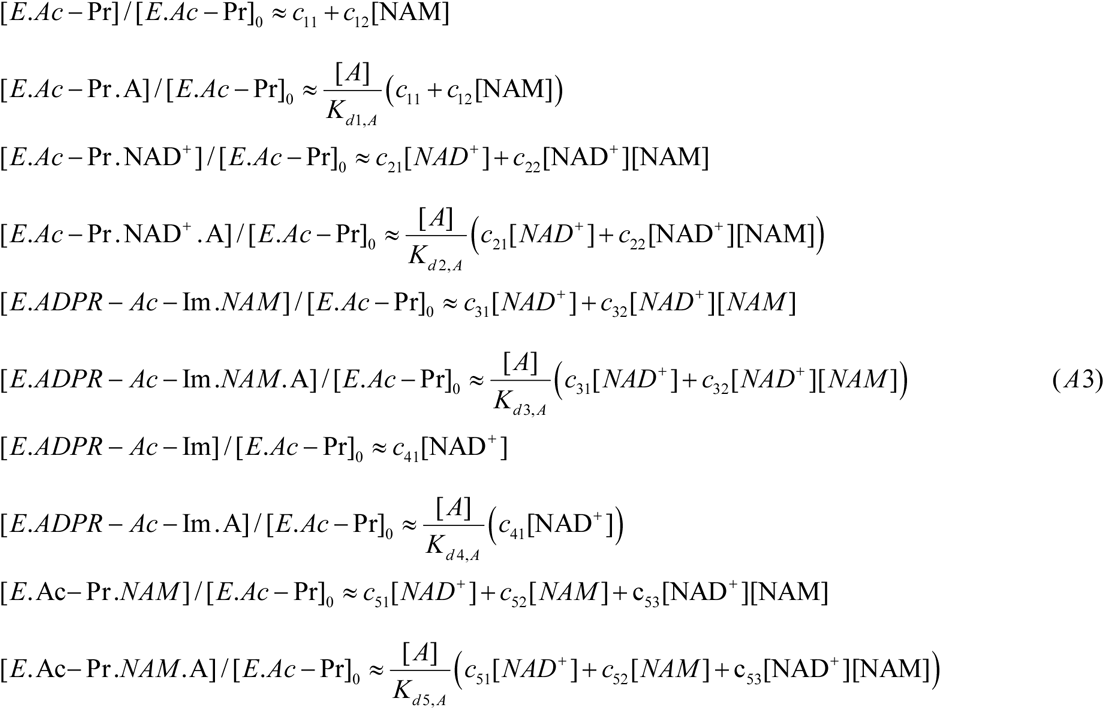

## References

[1] M. Kaeberlein, M. McVey, L. Guarente, The SIR2/3/4 complex and SIR2 alone promote longevity in Saccharomyces cerevisiae by two different mechanisms, Genes & Development, 13 (1999) 2570-2580.

[2] B.M. Hirsch, W. Zheng, Sirtuin mechanism and inhibition: explored with N(epsilon)-acetyl-lysine analogs, Molecular bioSystems, 7 (2011) 16-28.

[3] Y. Cen, Sirtuins inhibitors: the approach to affinity and selectivity, Biochimica et biophysica acta, 1804 (2010) 1635-1644.

[4] A.A. Sauve, Sirtuin chemical mechanisms, Biochimica Et Biophysica Acta-Proteins and Proteomics, 1804 (2010) 1591-1603.

[5] P. Hu, S. Wang, Y. Zhang, Highly dissociative and concerted mechanism for the nicotinamide cleavage reaction in Sir2Tm enzyme suggested by ab initio QM/MM molecular dynamics simulations, Journal of the American Chemical Society, 130 (2008) 16721-16728.

[6] Y. Zhou, H. Zhang, B. He, J. Du, H. Lin, R.A. Cerione, Q. Hao, The bicyclic intermediate structure provides insights into the desuccinylation mechanism of human sirtuin 5 (SIRT5), The Journal of biological chemistry, 287 (2012) 28307-28314.

[7] E.M. Mercken, S.J. Mitchell, A. Martin-Montalvo, R.K. Minor, M. Almeida, A.P. Gomes, M. Scheibye-Knudsen, H.H. Palacios, J.J. Licata, Y. Zhang, K.G. Becker, H. Khraiwesh, J.A. Gonzalez-Reyes, J.M. Villalba, J.A. Baur, P. Elliott, C. Westphal, G.P. Vlasuk, J.L. Ellis, D.A. Sinclair, M. Bernier, R. de Cabo, SRT2104 extends survival of male mice on a standard diet and preserves bone and muscle mass, Aging Cell, 13 (2014) 787-796.

[8] S.J. Mitchell, A. Martin-Montalvo, E.M. Mercken, H.H. Palacios, T.M. Ward, G. Abulwerdi, R.K. Minor, G.P. Vlasuk, J.L. Ellis, D.A. Sinclair, J. Dawson, D.B. Allison, Y. Zhang, K.G. Becker, M. Bernier, R. de Cabo, The SIRT1 activator SRT1720 extends lifespan and improves health of mice fed a standard diet, Cell reports, 6 (2014) 836-843.

[9] J.A. Zorn, J.A. Wells, Turning enzymes ON with small molecules, Nature chemical biology, 6 (2010) 179-188.

[10] D.A. Sinclair, L. Guarente, Small-Molecule Allosteric Activators of Sirtuins, Annual review of pharmacology and toxicology, 54 (2014) 363-380.

[11] B.P. Hubbard, A.P. Gomes, H. Dai, J. Li, A.W. Case, T. Considine, T.V. Riera, J.E. Lee, E.S. Yen, D.W. Lamming, B.L. Pentelute, E.R. Schuman, L.A. Stevens, A.J.Y. Ling, S.M. Armour, S. Michan, H.Z. Zhao, Y. Jiang, S.M. Sweitzer, C.A. Blum, J.S. Disch, P.Y. Ng, K.T. Howitz, A.P. Rolo, Y. Hamuro, J. Moss, R.B. Perni, J.L. Ellis, G.P. Vlasuk, D.A. Sinclair, Evidence for a Common Mechanism of SIRT1 Regulation by Allosteric Activators, Science, 339 (2013) 1216-1219.

[12] H. Dai, A.W. Case, T.V. Riera, T. Considine, J.E. Lee, Y. Hamuro, H.Z. Zhao, Y. Jiang, S.M. Sweitzer, B. Pietrak, B. Schwartz, C.A. Blum, J.S. Disch, R. Caldwell, B. Szczepankiewicz, C. Oalmann, P.Y. Ng, B.H. White, R. Casaubon, R. Narayan, K. Koppetsch, F. Bourbonais, B. Wu, J.F. Wang, D.M. Qian, F. Jiang, C. Mao, M.H. Wang, E. Hu, J.C. Wu, R.B. Perni, G.P. Vlasuk, J.L. Ellis, Crystallographic structure of a small molecule SIRT1 activator-enzyme complex, Nature Communications, 6 (2015) 7645.

[13] B.J. North, M.A. Rosenberg, K.B. Jeganathan, A.V. Hafner, S. Michan, J. Dai, D.J. Baker, Y.N. Cen, L.E. Wu, A.A. Sauve, J.M. van Deursen, A. Rosenzweig, D.A. Sinclair, SIRT2 induces the checkpoint kinase BubR1 to increase lifespan, Embo Journal, 33 (2014) 1438-1453.

[14] K. Brown, S. Xie, X. Qiu, M. Mohrin, J. Shin, Y. Liu, D. Zhang, D.T. Scadden, D. Chen, SIRT3 reverses aging-associated degeneration, Cell reports, 3 (2013) 319-327.

[15] Y. Kanfi, S. Naiman, G. Amir, V. Peshti, G. Zinman, L. Nahum, Z. Bar-Joseph, H.Y. Cohen, The sirtuin SIRT6 regulates lifespan in male mice, Nature, 483 (2012) 218-221.

[16] M. Gertz, F. Fischer, G.T.T. Nguyen, M. Lakshminarasimhan, M. Schutkowski, M. Weyand, C. Steegborn, Ex-527 inhibits Sirtuins by exploiting their unique NAD(+)-dependent deacetylation mechanism, Proceedings of the National Academy of Sciences of the United States of America, 110 (2013) E2772-E2781.

[17] T. Rumpf, M. Schiedel, B. Karaman, C. Roessler, B.J. North, A. Lehotzky, J. Olah, K.I. Ladwein, K. Schmidtkunz, M. Gajer, M. Pannek, C. Steegborn, D.A. Sinclair, S. Gerhardt, J. Ovadi, M. Schutkowski, W. Sippl, O. Einsle, M. Jung, Selective Sirt2 inhibition by ligand-induced rearrangement of the active site, Nature Communications, 6 (2015) 6263.

[18] A.P. Gomes, N.L. Price, A.J. Ling, J.J. Moslehi, M.K. Montgomery, L. Rajman, J.P. White, J.S. Teodoro, C.D. Wrann, B.P. Hubbard, E.M. Mercken, C.M. Palmeira, R. de Cabo, A.P. Rolo, N. Turner, E.L. Bell, D.A. Sinclair, Declining NAD(+) induces a pseudohypoxic state disrupting nuclear-mitochondrial communication during aging, Cell, 155 (2013) 1624-1638.

[19] V.B. Pillai, S. Samant, N.R. Sundaresan, H. Raghuraman, G. Kim, M.Y. Bonner, J.L. Arbiser, D.I. Walker, D.P. Jones, D. Gius, M.P. Gupta, Honokiol blocks and reverses cardiac hypertrophy in mice by activating mitochondrial Sirt3, Nature Communications, 6 (2015) 6656.

[20] A. Mai, S. Valente, S. Meade, V. Carafa, M. Tardugno, A. Nebbioso, A. Galmozzi, N. Mitro, E. De Fabiani, L. Altucci, A. Kazantsev, Study of 1,4-Dihydropyridine Structural Scaffold: Discovery of Novel Sirtuin Activators and Inhibitors, Journal of Medicinal Chemistry, 52 (2009) 5496-5504.

[21] S. Valente, P. Mellini, F. Spallotta, V. Carafa, A. Nebbioso, L. Polletta, I. Carnevale, S. Saladini, D. Trisciuoglio, C. Gabellini, M. Tardugno, C. Zwergel, C. Cencioni, S. Atlante, S. Moniot, C. Steegborn, R. Budriesi, M. Tafani, D. Del Bufalo, L. Altucci, C. Gaetano, A. Mai, 1,4-Dihydropyridines Active on the SIRT1/AMPK Pathway Ameliorate Skin Repair and Mitochondrial Function and Exhibit Inhibition of Proliferation in Cancer Cells, Journal of Medicinal Chemistry, 59 (2016) 1471-1491.

[22] W. You, D. Rotili, T.M. Li, C. Kambach, M. Meleshin, M. Schutkowski, K.F. Chua, A. Mai, C. Steegborn, Structural Basis of Sirtuin 6 Activation by Synthetic Small Molecules, Angewandte Chemie (International ed. in English), 56 (2017) 1007-1011.

[23] J.L. Feldman, K.E. Dittenhafer-Reed, N. Kudo, J.N. Thelen, A. Ito, M. Yoshida, J.M. Denu, Kinetic and Structural Basis for Acyl-Group Selectivity and NAD+ Dependence in Sirtuin-Catalyzed Deacylation, Biochemistry, 54 (2015) 3037-3050.

[24] B.C. Smith, J.M. Denu, Sir2 deacetylases exhibit nucleophilic participation of acetyl-lysine in NAD+ cleavage, Journal of the American Chemical Society, 129 (2007) 5802-5803.

[25] J.L. Avalos, J.D. Boeke, C. Wolberger, Structural basis for the mechanism and regulation of Sir2 enzymes, Molecular Cell, 13 (2004) 639-648.

[26] Z. Liang, T. Shi, S. Ouyang, H. Li, K. Yu, W. Zhu, C. Luo, H. Jiang, Investigation of the catalytic mechanism of Sir2 enzyme with QM/MM approach: SN1 vs SN2?, The journal of physical chemistry. B, 114 (2010) 11927-11933.

[27] Y. Shi, Y. Zhou, S. Wang, Y. Zhang, Sirtuin Deacetylation Mechanism and Catalytic Role of the Dynamic Cofactor Binding Loop, The Journal of Physical Chemistry Letters, 4 (2013) 491-495.

[28] J.L. Avalos, K.M. Bever, C. Wolberger, Mechanism of sirtuin inhibition by nicotinamide: Altering the NAD(+) cosubstrate specificity of a Sir2 enzyme, Molecular Cell, 17 (2005) 855-868.

[29] X. Guan, P. Lin, E. Knoll, R. Chakrabarti, Mechanism of inhibition of the human sirtuin enzyme SIRT3 by nicotinamide: computational and experimental studies, PLoS One, 9 (2014) e107729.

[30] Y. Cen, A.A. Sauve, Transition state of ADP-ribosylation of acetyllysine catalyzed by Archaeoglobus fulgidus Sir2 determined by kinetic isotope effects and computational approaches, Journal of the American Chemical Society, 132 (2010) 12286-12298.

[31] A.A. Sauve, R.D. Moir, V.L. Schramm, I.M. Willis, Chemical activation of Sir2-dependent silencing by relief of nicotinamide inhibition, Molecular Cell, 17 (2005) 595-601.

[32] W.F. Hawse, K.G. Hoff, D.G. Fatkins, A. Daines, O.V. Zubkova, V.L. Schramm, W. Zheng, C. Wolberger, Structural insights into intermediate steps in the Sir2 deacetylation reaction, Structure, 16 (2008) 1368-1377.

[33] C.M. Armstrong, M. Kaeberlein, S.I. Imai, L. Guarente, Mutations in Saccharomyces cerevisiae gene SIR2 can have differential effects on in vivo silencing phenotypes and in vitro histone deacetylation activity, Molecular Biology of the Cell, 13 (2002) 1427-1438.

[34] B.G. Szczepankiewicz, H. Dai, K.J. Koppetsch, D. Qian, F. Jiang, C. Mao, R.B. Perni, Synthesis of carba-NAD and the structures of its ternary complexes with SIRT3 and SIRT5, The Journal of organic chemistry, 77 (2012) 7319-7329.

[35] K.H. Zhao, R. Harshaw, X.M. Chai, R. Marmorstein, Structural basis for nicotinamide cleavage and ADP-ribose transfer by NAD(+)-dependent Sir2 histone/protein deacetylases, Proceedings of the National Academy of Sciences of the United States of America, 101 (2004) 8563-8568.

[36] M. Pacholec, J.E. Bleasdale, B. Chrunyk, D. Cunningham, D. Flynn, R.S. Garofalo, D. Griffith, M. Griffor, P. Loulakis, B. Pabst, X. Qiu, B. Stockman, V. Thanabal, A. Varghese, J. Ward, J. Withka, K. Ahn, SRT1720, SRT2183, SRT1460, and resveratrol are not direct activators of SIRT1, The Journal of biological chemistry, 285 (2010) 8340-8351.

[37] S. Imai, A possibility of nutriceuticals as an anti-aging intervention: activation of sirtuins by promoting mammalian NAD biosynthesis, Pharmacological research, 62 (2010) 42-47.

[38] G. Wang, T. Han, D. Nijhawan, P. Theodoropoulos, J. Naidoo, S. Yadavalli, H. Mirzaei, A.A. Pieper, J.M. Ready, S.L. McKnight, P7C3 neuroprotective chemicals function by activating the rate-limiting enzyme in NAD salvage, Cell, 158 (2014) 1324-1334.

[39] W.F. Hawse, C. Wolberger, Structure-based Mechanism of ADP-ribosylation by Sirtuins, Journal of Biological Chemistry, 284 (2009) 33654-33661.

[40] R. Chakrabarti, A.M. Klibanov, R.A. Friesner, Sequence optimization and designability of enzyme active sites, Proceedings of the National Academy of Sciences of the United States of America, 102 (2005) 12035-12040.

